# Hypometabolism to survive the long polar night in the diatom *Fragilariopsis cylindrus*

**DOI:** 10.1101/2023.01.14.524047

**Authors:** Nathalie Joli, Lorenzo Concia, Karel Mocaer, Julie Guterman, Juliette Laude, Sebastien Guerin, Theo Sciandra, Flavienne Bruyant, Ouardia Ait-Mohamed, Marine Beguin, Marie-Helene Forget, Clara Bourbousse, Thomas Lacour, Benjamin Bailleul, Jean-Eric Tremblay, Douglas Campbell, Johan Lavaud, Yannick Schwab, Marcel Babin, Chris Bowler

**Affiliations:** Institut de Biologie de l’École Normale Supérieure (IBENS), École Normale Supérieure, CNRS, INSERM, PSL Université Paris, 75005 Paris, France; Cell Biology and Biophysics Unit, European Molecular Biology Laboratory (EMBL), 69117 Heidelberg, Germany; Takuvik International Research Laboratory, Université Laval (Canada) & CNRS (France), Département de Biologie and Québec-Océan, Université Laval, Pavillon Alexandre-Vachon 1045, avenue de la Médecine, Local 2078, G1V 0A6, Canada; Laboratoire PHYSiologie des micro ALGues (PDG-ODE-PHYTOX-PHYSALG), Centre Atlantique, Rue de l’Ile d’Yeu, BP 21105, 44 311 Nantes, Cedex 03, France; Laboratory of Chloroplast Biology and Light Sensing in Microalgae, Institut de Biologie Physico Chimique, CNRS, Sorbonne Université, Paris 75005, France; Département de Biologie, Université Laval, Québec, QC, Canada; Mount Allison University, Sackville, New Brunswick, Canada; LIENSs ‘Littoral, Environnement et Sociétés’ (UMRi7266), CNRS/Université de La Rochelle, Institut du Littoral et de l’Environnement, La Rochelle, France

## Abstract

Diatoms, the major eukaryotic phytoplankton in polar regions, are essential to sustain Arctic and Antarctic ecosystems. As such, it is fundamental to understand the physiological mechanisms and associated molecular basis of their resilience to the long polar night. Here, we report an integrative approach revealing that in prolonged darkness, diatom cells enter a state of quiescence associated with reduced metabolic and transcriptional activity during which no cell division occurs. We propose that minimal energy is provided by respiration and degradation of protein, carbohydrate, and lipid stores and that homeostasis is maintained by autophagy in prolonged darkness. We also report internal structural changes that manifest the morphological acclimation of cells to darkness. Our results further indicate that immediately following a return to light, diatom cells are able to use photoprotective mechanisms and rapidly resume photosynthesis. Cell division resumed rates similar to those before darkness. Our study demonstrates the remarkable robustness of polar diatoms to prolonged darkness at low temperatures.

**Graphical abstract:** 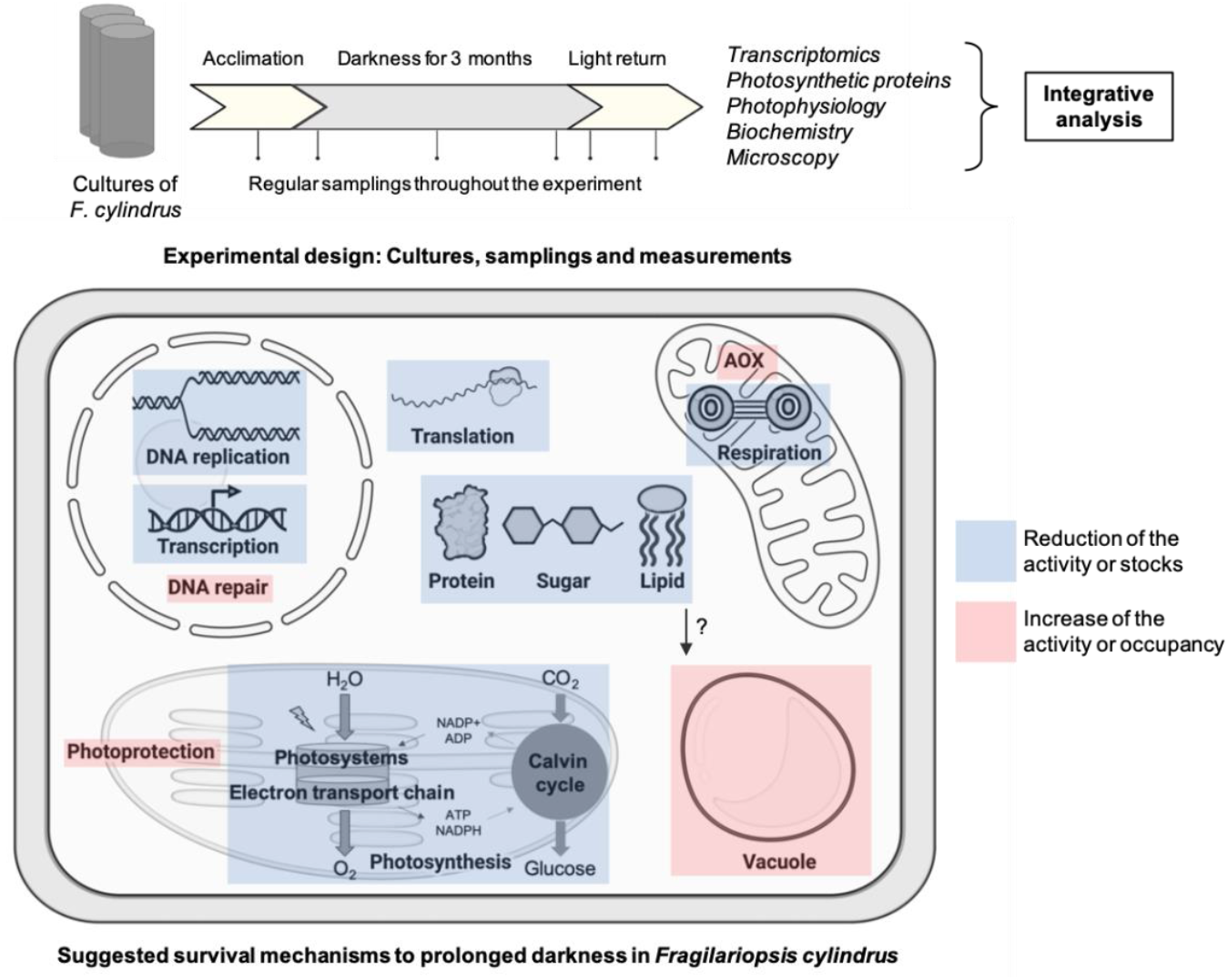

**Teaser:** To survive the long winter, polar diatoms slow down metabolism and express genes to assure survival following return to light.

## Introduction

Polar regions are characterized by year-round cold, continuous light around summer solstice and continuous darkness around winter solstice. Species, ranging from plankton to mammals and birds, adapted to polar conditions can occur in large numbers of individuals (*1*, *2*) but need to survive through winter darkness, when energy is limited. The survival of polar phytoplankton, which are the basis of food chains in the Arctic and Antarctic, over winter is little understood. Light is variable at polar latitudes with up to several months of continuous darkness or light, separated by transition periods where photoperiod changes can reach 35 minutes (min) per day (d), and by variable local light conditions due to ice and snow cover. The ability of phytoplankton to survive during the polar night, despite the prolonged quasi-absence of light, has been reported for over 30 years (*3*, *4*), and the size of the surviving population of over-wintering phytoplankton is thought to impact the level of primary production once light is again available after annual sea ice breakup (*5*). The effects of global warming are particularly acute in the polar regions, so understanding the resilience mechanisms of the organisms there and their tolerance to environmental change is now of pressing importance, as they transition towards warmer temperatures while still enduring extreme variations in seasonal light.

High latitude phytoplankton biomass is dominated by diatoms (*6*), unicellular eukaryotes with an external cell wall made of silica, known as frustule. Diatoms are photosynthetic Stramenopiles, with chloroplasts derived via secondary endosymbiosis contributing to a combination of host- and endosymbiont-derived genes in their nuclear genome (*7*). Diatoms play key roles in the global carbon and silica cycles in the ocean (*8*), accounting for about half of marine primary production (*9*). The resilience of diatoms to rollercoaster fluctuations in environmental conditions, such as those prevailing in polar oceans, may be one of the keys to their ecological success. They can notably cope with highly variable light conditions.

Diatoms have evolved efficient mechanisms to cope with rapid and extreme environmental fluctuations. Their photosynthetic pigments differ significantly from those of green algae and plants, including chlorophyll c (Chl c), fucoxanthin (Fx), and the photoprotective pigments diadinoxanthin (DD) and diatoxanthin (DT) (*10*). In addition, diatoms possess gene subfamilies encoding light harvesting complex proteins (LHCx and LHCz), which are believed to enhance photosynthetic light capture under variable environmental conditions (*11*), and cryptochromes that have a dual function in photoperception and DNA repair (*12*). Their success at high latitudes and in sea ice habitats (*2*, *15*) indicates that diatoms have strategies to survive several months of darkness. Although heterotrophic nutrition has been documented as a means of survival in the dark (*13*, *14*), the lack of references to autotrophic diatoms capable of active growth in the dark leads us to consider this mode of nutrition only secondarily in these organisms. Furthermore, a recent functional classification of eukaryotic unicellular plankton suggests that diatoms are the only group with an obligate photoautotrophic lifestyle within the Stramenopiles (*15*). Some studies have focused on the survival metabolism of diatoms during short periods of darkness (weeks) (*16*–*19*) while others have shown that some diatoms can survive much longer in darkness without the addition of organic sources (*5*, *20*, *4*, *21*).

Under adverse conditions, microalgae including diatoms re-organize membrane and plastid lipids into lipid droplets (LD) comprised of storage lipids to support survival. While fatty acid synthesis is strongly suppressed at the transcript level, triacylglycerol (TAG) accumulation is observed (*22*), raising the hypothesis that this is a consequence of C reallocation. The major carbohydrate food reserve of diatoms is chrysolaminarin (CHY), a β-1,3-glucan polysaccharide stored in intracellular vacuoles (*23*). In *Thalassiosira pseudonana* and *Phaeodactylum tricornutum*, CHY degradation genes are induced under phosphate-limited conditions (*24*). In addition, under nitrogen deprivation, it was observed that diatoms do not proliferate but are able to resume growth once favorable conditions return (*25*), a characteristic of quiescent cells (*26*).

*Fragilariopsis* is considered one of the most abundant and diverse genera in the global ocean and *F. cylindrus* is believed to be among the most successful in the Southern Ocean (*27*). Having adapted to low light conditions such as is found under sea ice, this species represents a valuable organism to explore life in the polar environment, complemented by the availability of a reference genome (*19*). While dark survival of *F. cylindrus* has received broad attention, there is typically an uncoupling between the methods used. For example, some authors (*28*) monitored multiple physiological and metabolic traits of *F. cylindrus* in the dark while others (*19*) characterized gene expression without physiological analyses, and light recovery was not studied. Recently, a respiratory metabolism of dark-acclimated cells using mostly proteomic profiling has been proposed (*29*). Consequently, we lack an integrated understanding of how this diatom survives the long polar night.

Here we developed an experimental setup using a large culture volume (80 L) that allows for a combination of microscopic, transcriptomic and biochemical approaches to assess survival strategies during an extended period of darkness and subsequent re-exposure to light. We show that, in darkness, cells enter a quiescent state during which no cell division occurs, associated with reduced transcriptional and translational activities. During this period, the cell provides minimal energy supply by respiration via the alternative oxidase pathway, degradation of stored carbohydrates and lipids, and ensures safe resumption of metabolic activity in the light by DNA repair and photoprotection. Sufficient photosynthetic pigments are retained and photosystems remain in a functional state to convert light energy into photochemistry immediately when light returns. We also describe internal structural changes, such as vacuole formation, and propose potential roles in the dark survival strategy. Although the sudden return of light is associated with the death of about one-third of the population, the majority of cells were able to use photoprotective mechanisms, resume photosynthesis, and divide within a few days. Our study demonstrates the resilience of *F. cylindrus* to the long polar night.

## Results

### Cellular bioenergetics during the initial acclimation phase in full light, prolonged darkness, and final return to light

Three biological replicates of *F. cylindrus* cultures acclimated to continuous light were subjected to darkness for a period of 86d followed by exposure to continuous light for 7d. Numerous parameters were measured, summarized in **<u>Table S1</u>**. We firstly explored bulk and within-cell biochemical and photophysiological changes. Except for the increase in cell abundance (up to 30%) observed during the first 3d, likely resulting from the completion of the last division, *F. cylindrus* stops cell division in the dark (**Figure 1A, <u>S1A</u>)**. Viability as determined by cytometry (SYTOX) remained high (99%) during the dark period. Following light return, the increase in cell number from day 3 onward is consistent with the time required for division, in addition to the time required to significantly exceed the 30% mortality rate that occurs from 30 min up to 3d (**<u>Figure S1B</u>**). This ability to rapidly resume growth within a few days of the return of light after a prolonged period of darkness seems to be a common characteristic of polar diatoms (*21*).

**Figure 1:**
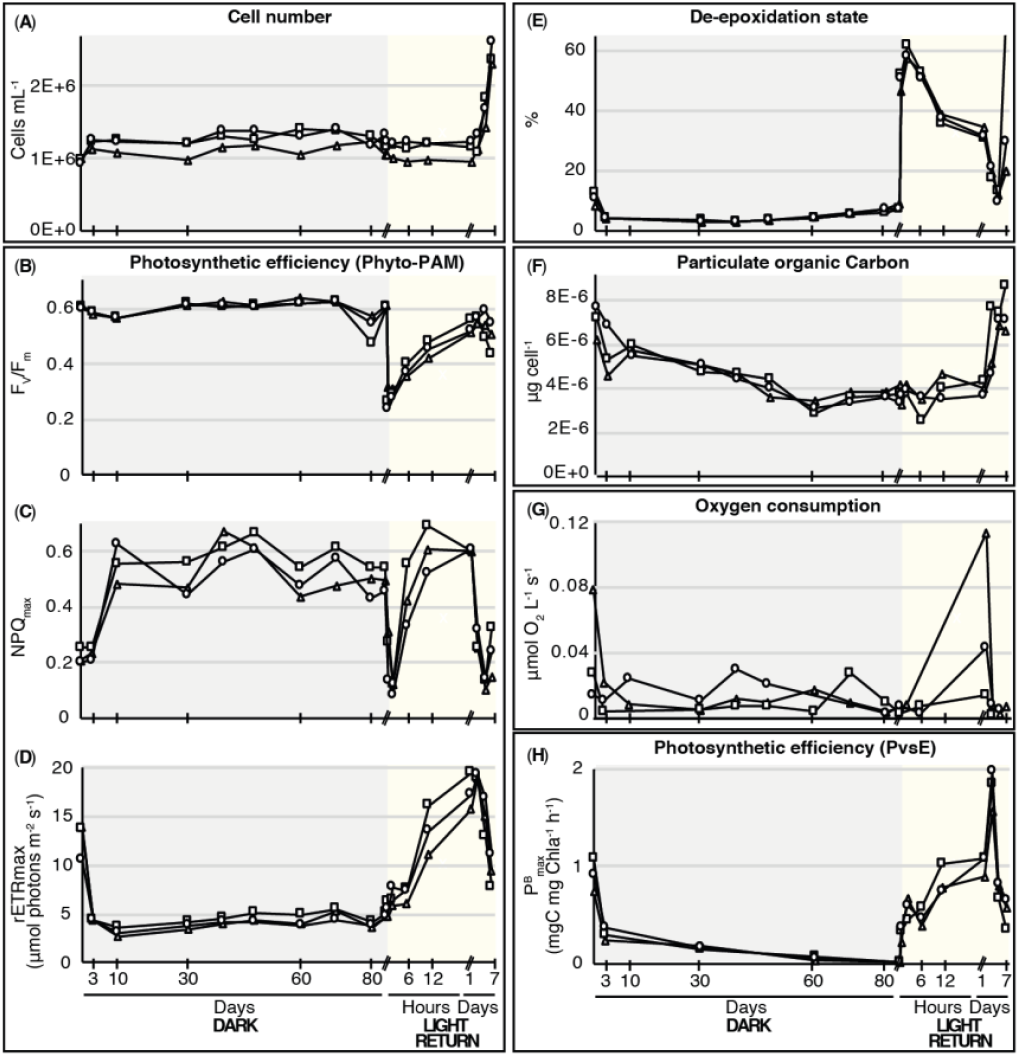
Cellular bioenergetics. Tracking physiological changes from the acclimation phase in full light, in prolonged darkness (grey area of the graphs), and upon return to light (yellow area of the graphs) (**A**) Number of cells per mL (**B**) Maximum quantum yield of PSII (F_V_/F_m_) (**C**) Non-Photochemical Quenching (NPQ_max_) (**D**) Maximum relative photosynthetic electron transport rate (rETR_max_, μmol photons m^-2^ s^-1^) (**E**) De-epoxidation state (DES, %). (**F**) Carbon concentration (μg cell^-1^) (**G**) Oxygen consumption (μmol O_2_ L^-1^ s^-1^) (**H**) Maximum specific carbon fixation rate in mgC mg Chla^-1^ h^-1^ (P^B^_max_). The three biological replicates are represented with triangles (Cylinder 1), squares (Cylinder 2) and circles (Cylinder 3). X axis shows the time points sampled with a broken-scale in hours (h) or days (d). See Table S1 and Figure S1 for all the time points sampled per parameter.

Bulk particulate carbon and nitrogen concentrations, together with photosynthetic pigments per cell (Chl a, Chl c2 and Fx), decrease by half during the dark period, indicating the potential consumption of internal compounds (**Figure 1F, <u>S1D, S1F</u>**). Lipid content as estimated by cytometry (BODIPY) also gradually decreased in prolonged darkness (**<u>Figure S1B</u>**). Although the measured values fluctuated during the dark period, we observed a general decrease in oxygen consumption after 3 months of darkness, being 2 to 13 times lower than the value measured during the light acclimation period prior to the dark period, depending on the cylinder considered (**Figure 1G, <u>S1E</u>**). A respiration maximum was observed 1d after light return. Within 2 hours (h) of the return to light, the de-epoxidation state of xanthophyll cycle pigments (DES = DT/(DD+DT) x 100) reached a maximum of 60%, illustrating the induction of photoprotection through non-photochemical quenching (**Figure 1C, 1E, <u>S1G</u>**). While lipid stocks are replenished within the first 2d of light return, particulate carbon and nitrogen, as well as photosynthetic pigment stocks, were restored to baseline after 5d of light return.

The maintenance of a high photochemical efficiency of photosystem II F_V_/F_m_ throughout the experiment along with the slight decrease in the average absorption cross section σPSII (**Figure 1B, <u>S1H, S1I</u>**) indicate the conservation of the ability to convert light energy by photochemistry in the dark (*18*, *21*, *28*). However, we observed a drastic decrease in capacity for photosynthetic electron transfer (rETR_max_), in carbon fixation (P^B^_max_) and in photosynthetic protein contents (RuBisCo large subunit protein RbcL and the photosystem II (PSII) protein D1 PsbA) within the first 3d of darkness (**Figure 1D, 1H, <u>S1H, S1J, S1K</u>**). All the previous mentioned parameters returned to baseline within 1d of light return.

The cell appears to be in a state of quiescence, temporarily exiting the cell cycle but still retaining the ability to reinitiate cell division (*41*, *42*). In agreement with previous studies in polar diatoms, the decrease in photochemical potential observed here reflects the acclimation of *F. cylindrus* to prolonged darkness (*4*, *20*, *21*, *28*).

### Morphological changes of F. cylindrus in continuous light and after prolonged darkness

We then assessed the morphological responses of *F. cylindrus* to prolonged darkness by microscopy. Scanning electron microscopy (SEM), transmission electron microscopy (TEM) and focused ion beam-scanning electron microscopy (FIB-SEM) were performed on light- and dark-acclimated cells (2 months). SEM did not reveal any external differences, indicating that *F. cylindrus* does not form spores in the dark (**Figure 2A**). FIB-SEM and TEM revealed significant changes in cell constitution, including the presence of a newly formed vacuole in prolonged darkness (**Figure 2B, 2C, 3A**).

**Figure 2:**
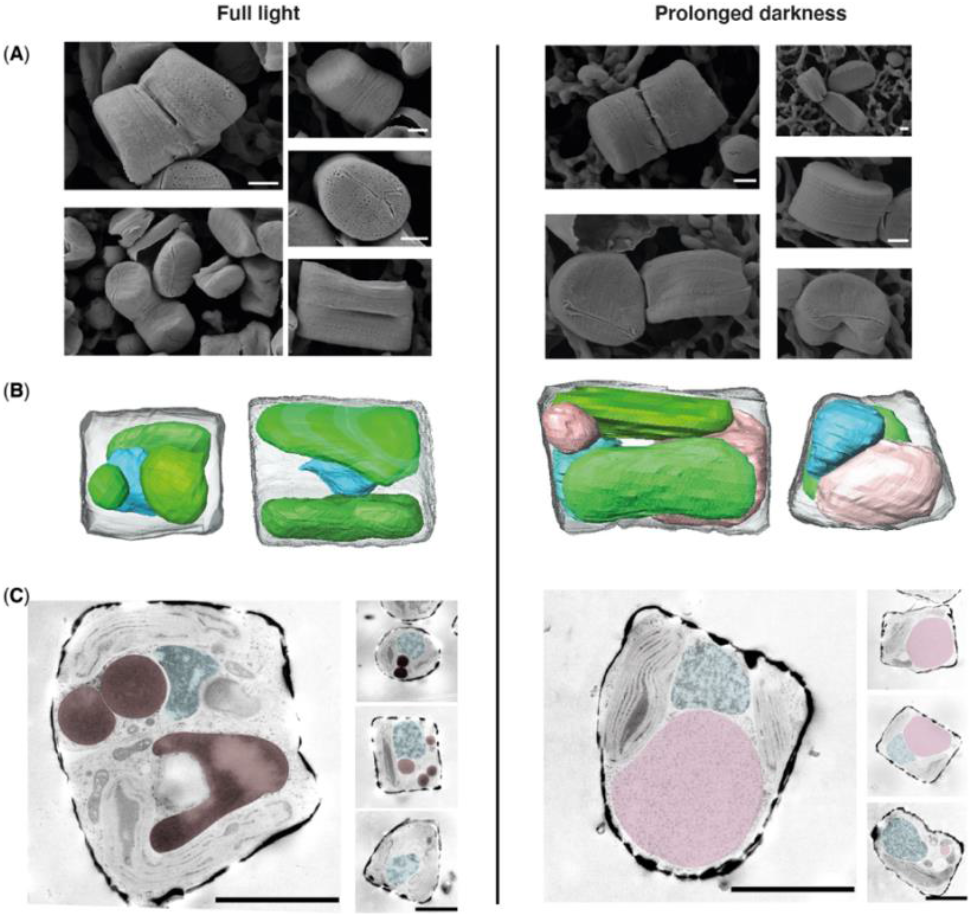
Morphological changes of *F. cylindrus* in continuous light (left panels) and after prolonged darkness (right panels). (**A**) Outer cell morphology observed by SEM. (**B**) 3D cell reconstruction from FIB-SEM. Chloroplasts are in green, the nucleus in light blue and the vacuole in pink (See supplemental videos). (**C**) Inner cell morphology observed by SEM. Lipids are in brown, the nucleus in light blue and the vacuole in pink. For each image, the scale bar represents 1μm.

**Figure 3:**
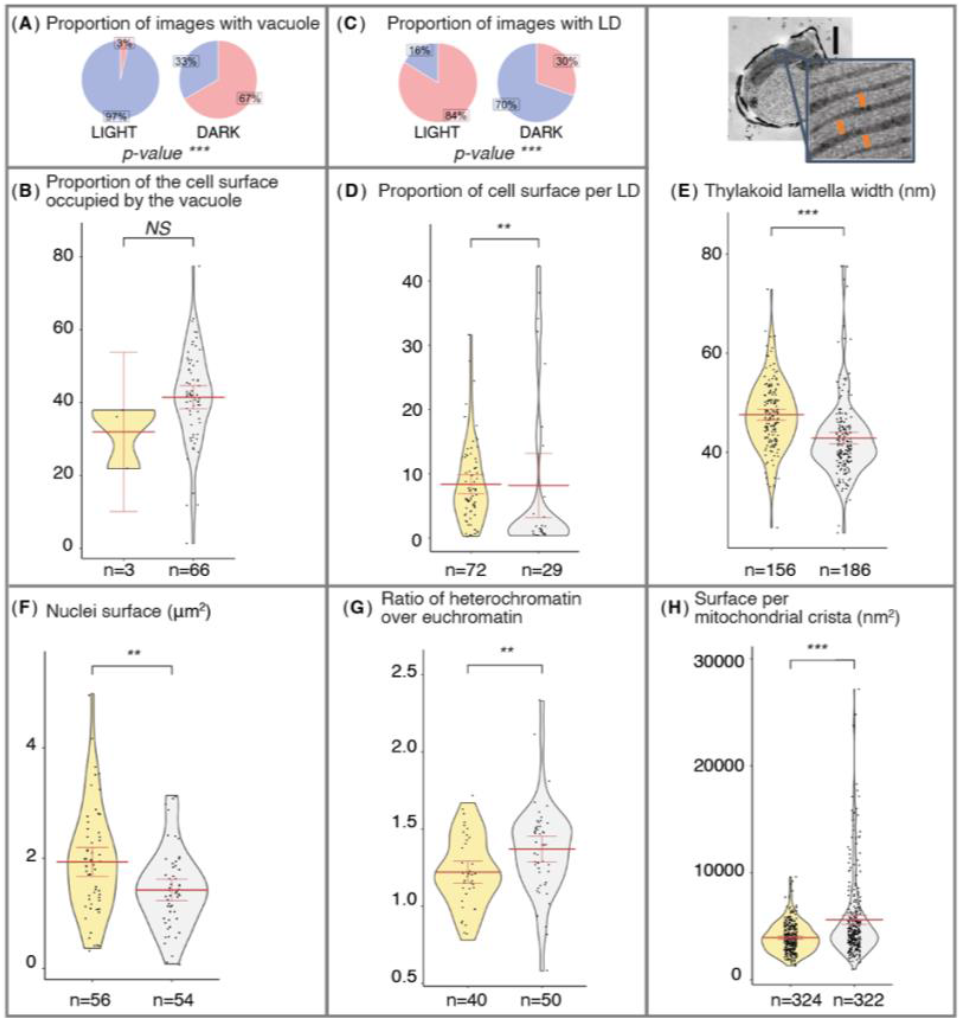
TEM image analyses of *F. cylindrus* cells in light (yellow plots) and in dark (grey plots) conditions show distinctive structural features. (**A**) Diagram comparing the proportion of light/dark cells that do not contain a vacuole (blue) to those that do (red). (**B**) Violin plot displaying light/dark cells proportion occupied by the vacuole. (**C**) Diagram comparing the proportion of light/dark cells that do not contain lipid droplets (LD) (blue) to those that do (red). (**D**) Violin plot depicting light/dark cells proportion occupied by lipid droplets (LD). (**E**) Images of a dark cell with a zoom on the thylakoid width measurement. Violin plots display the thylakoid lamellae width in nm within light/dark cell’s chloroplasts. Scale bar represents 500nm. (**F**) Violin plot comparing the light/dark cell’s nucleus areas. (**G**) Violin plot comparing the heterochromatin over euchromatin ratio of the light/dark cells. (**H**) Violin plot of the area in nm^2^ per mitochondrial cristae in light/dark cells. For all violin plots, the mean is shown by the bold horizontal bar; its 95% confidence interval is between the two error bars. n is the number of images used for each plot. Result of mean comparison is represented by its p-value above the plots. Not significant (NS), asterisk (*) is p-value (p) of 1% ≤ p < 5%; ** is 0.1% ≤ p < 1% and *** is p < 0.1%.

FIB-SEM results take into account the total volume of each cell (up to 5 cells). TEM images, on the other hand, reflect the proportions of a given organelle or feature from a thin cell section and measurements from these images must therefore be taken with caution despite the large number of cells considered (up to 99 cells). The large vacuole occupied 25% of the cell volume estimated by FIB-SEM, or 40% of the projected area estimated by TEM, while the cell area appears to remain unchanged according to TEM and FIB-SEM (**Figure 3B, <u>S2C, S3F</u>**). In agreement with cell counts, TEM analysis showed significantly fewer dark-acclimated dividing-cells (p < 0.1%) (**<u>Figure S3E</u>**). TEM image analysis confirms the significant decrease in the number and size of dense-core vesicles (0.1% ≤ p < 1%) that we consider to be LD, stained by NileRed in confocal microscopy (**Figure 3C, 3D, <u>S4</u>**). We also observed morphological changes within the chloroplast and mitochondria, with a reduction in thylakoid lamella width and more swollen mitochondrial cristae after 2 months of darkness (p < 0.1%) (**Figure 3E, 3H**). We observed ovoid-shaped droplets filled with whitish material reminiscent of the feature observed in other diatoms by immunostaining with an anti-β-1,3-glucan antibody by TEM (*23*). These are likely to be CHY stocks and appear to be larger in some dark-acclimated cells (**<u>Figure S3C</u>**), suggesting that cells may accumulate large amounts of carbohydrate in the dark, as has been observed in stressed or quiescent cells (*30*, *31*). The nucleolus (0.1% ≤ p < 1%) was smaller in the dark (**<u>Figure S3I</u>**). Since this organelle is known to be the ribosome factory, which in turn is responsible for protein production, this observation could underpin the decrease in metabolic activity. In addition, a higher ratio of heterochromatin to euchromatin was found in the dark which, in conjunction with the smaller size of the nucleus (0.1% ≤ p < 1%), could mean that the DNA is more condensed and thus less accessible for gene transcription (**Figure 3F, 3G**).

### Enriched functions within temporal clusters of gene expression

To explore temporal changes in gene expression, we sequenced a total of 48 transcriptomes from cultures that span the time series from the full-light acclimation phase before darkness (T0LIGHT), through darkness to the light return phase (**<u>Table S1</u>**). Principal component analysis revealed a gradual transition, first from 6h to 6d of darkness (short-term darkness), and then to 1 and 2 months of darkness (long-term darkness). We noted a continuous change from 30 min to 2h of return to light (short-term response to light) to 6h to 1d of return to light (longer-term response). The return to the initial state of gene expression consistent with T0LIGHT was observed after 2d of light return (**Figure 4A**). Of note, similar results were obtained for the transcriptome sequencing of another set of samples that spent only one month in the dark (**<u>Figure S5</u>**). Using the Dynamic Regulatory Events Miner approach, we could identify 5 temporal macro trends (clusters) in gene expression within 14,853 genes (62% of the total number of genes) as compared to T0LIGHT. Cluster 1 and Cluster 4 are composed of genes that maintain their expression in the dark. Cluster 1 genes are upregulated upon return of light, while Cluster 4 genes are downregulated relative to T0LIGHT. Cluster 2, Cluster 3, and Cluster 5 are all representatives of genes that are downregulated in the dark. While the genes included in Cluster 2 show a gradual increase in expression from the early light return period and Cluster 3 only after 1d of light return, Cluster 5 contains genes that are strongly downregulated throughout the experiment. Clusters were then subdivided into subclusters (**Figure 4B, 4C, <u>Table S2</u>**).

**Figure 4:**
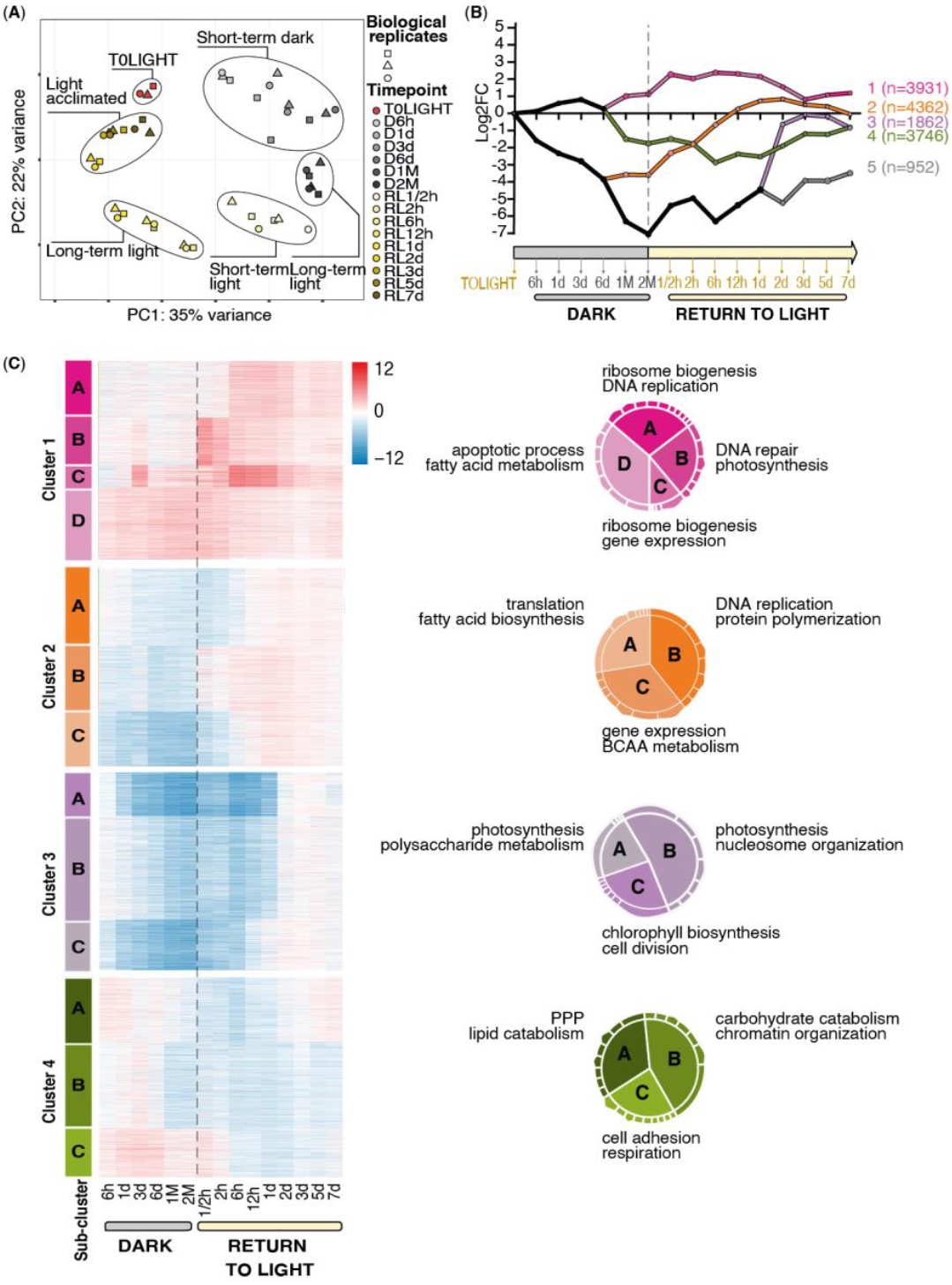
Gradual transition over time at the transcriptomic level in light/dark cells and enriched functions. (**A**) Principal component analysis of a total 48 light/dark transcriptomes from cultures in the acclimation phase in full light (T0LIGHT), in prolonged darkness (up to 2 months), and upon return to light (up to 7 days). Each dot represents one transcriptome and the shapes refer to one of the three biological replicates. (**B**) Dynamic Regulatory Events Miner analysis using the differentially expressed genes compared to the acclimation phase. The numbers on the right refer to the number of the cluster along with the number of genes within each. (**C**) Each of the clusters is divided into sub-clusters illustrated by letters on the left of each heatmap (A to C or A to D). Heatmaps show the log2FC of genes compared to the acclimation time point. Pie charts on the right of each heatmap indicate the gene ontology GO enrichment within each sub-cluster. Only two GOs per subcluster are shown (notched sectors). The full list of enriched GOs is given in **<u>Table S2</u>**.

The enriched functions within subclusters 1A to 1C suggest that the cell repairs DNA after 30min of light return, and that genes involved in gene expression and ribosome biogenesis were still activated after 3d of darkness, switched off in prolonged darkness, and turned on again after 6h of light return. Subcluster 1D, upregulated throughout the dark period and losing expression to return to baseline 3d after light return, is enriched in genes involved in apoptotic processes. Subclusters 2A and 2C have similar temporal trends, with downregulation in the dark and increased expression from 12h after light return, and are enriched in functions such as gene expression, translation and fatty acid synthesis. The enriched functions within subcluster 2B suggest that DNA replication restarts from 30 min after light return. The subclusters 3A to 3C are genes that recover the same expression level as T0LIGHT starting from 1-2 d after light return. They are enriched in genes involved in photosynthesis, chlorophyll synthesis and cell division. Subclusters 4A and 4B comprise genes that were overexpressed during the early darkness and are enriched in genes involved in chromatin organization, lipid and carbohydrate catabolism. Subcluster 4C contains genes that are induced throughout the dark period as compared to T0LIGHT, and is enriched in functions such as respiration (**Figure 4C**). The smallest Cluster 5 is made up of genes downregulated throughout the experiment compared to T0LIGHT and, unlike all other clusters, these genes do not return to the initial state after 7d of light return. The functions detected in this cluster include signal transduction, transcription and response to wounding (not illustrated here) (**<u>Table S2</u>**).

To detect fast variations in expression that could have been missed in the macrotrends analysis, we examined genes that were differentially expressed at single time points as compared to T0LIGHT. The list includes 3,619 genes (15% of total number of genes) among which one third are specific to 30 min light return. Among the enriched gene ontology GO categories, we found cell communication, mRNA and protein metabolism, and cytochrome complex assembly. A total of 194 genes were specifically differentially expressed after 3d of darkness; the enriched gene ontology GO functions include apoptosis (**<u>Figure S6</u>**). The complete list of genes is given in **<u>Table S3</u>**.

### Integrative view of cell metabolism in the dark and following light return

We then explored metabolic pathways of interest by relating patterns of gene expression (samples merged according to **Figure 4A**) with the corresponding biochemical results when available. The results are summarized in **Figure 5**. The timing of the transition from short-term to long-term dark response at the transcriptomic level is consistent with the stabilization of cell counts in the culture, along with the decrease in rETR_max_, P^B^_max_, and in the contents of the photosynthetic proteins RbcL and PsbA after several days in darkness (**Figure 1A, 1D, 1H, 4A, 5**).

**Figure 5:**
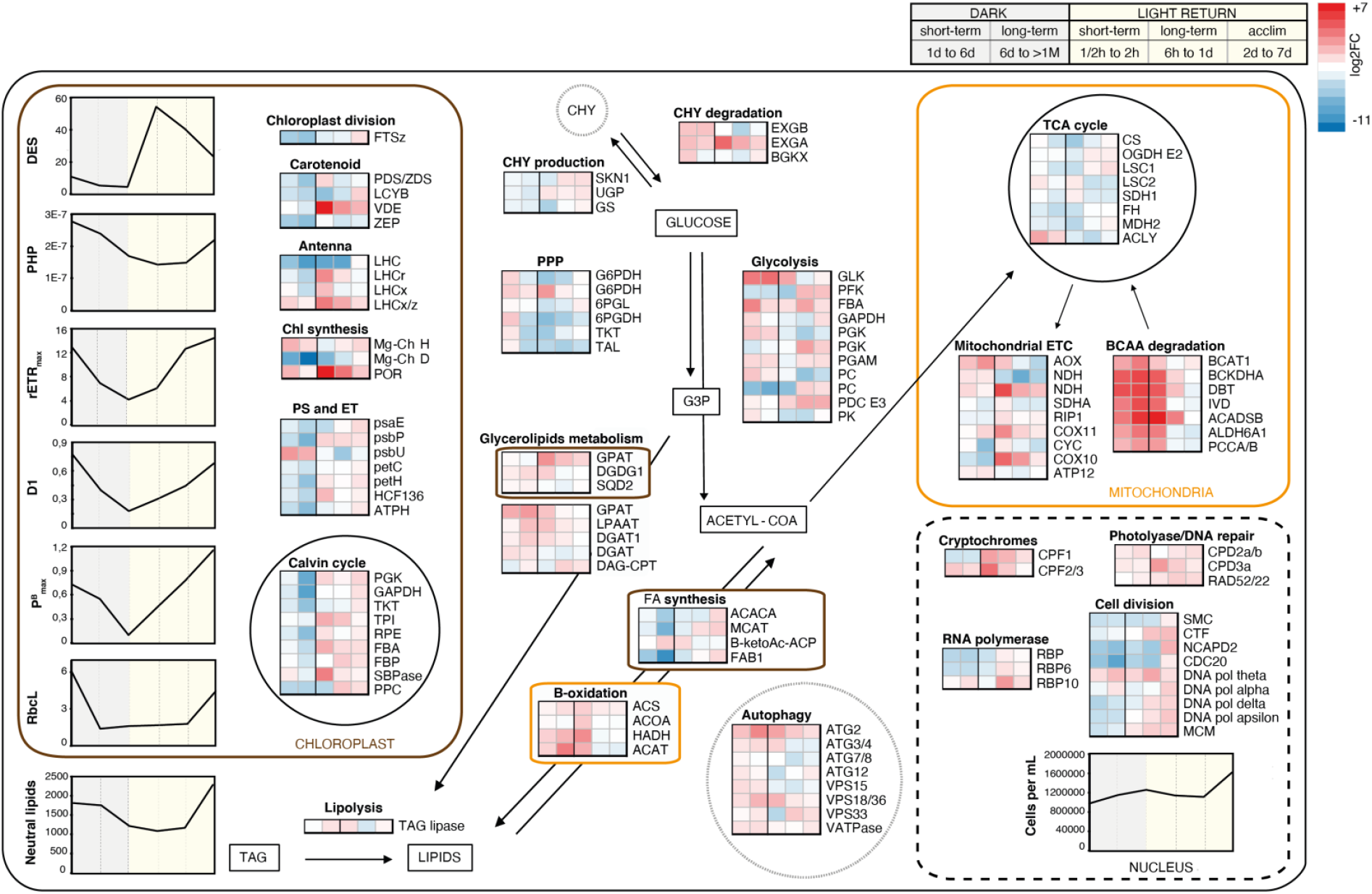
Cell schematic displaying average expression of selected genes of interest in *F. cylindrus*. Colored squares indicate the average expression change (log2FC) in each time window relative to the pre-dark acclimation phase. Red represents higher expression and blue represents lower expression. Short-term dark (1 to 6 days), long-term dark (>6 days to 2 months), short-term light return (30 minutes to 2 hours), long-term light return (6 hours to 1 day), and light acclimation (2 to 7 days) conditions are represented. Genes involved in metabolic pathways located in the chloroplast are represented inside a brown box, those in the mitochondria inside an orange box. Those in the nucleus are shown inside a black dashed box and those in vacuoles in grey dotted circles. Percentage de-epoxidation state (DES), sum of photosynthetic pigments (PHP = Chl a + Chl c + Fx) in μg cell^-1^, thylakoid membrane electron transport rate (rETR), maximum productivity (PBmax), neutral lipids expressed in relative fluorescence units as well as photosynthetic proteins are represented (RbcL and D1 in μg). The complete list of corresponding gene names with their acronyms, the number of genes considered and the subcellular prediction is given in **<u>Table S4</u>.** Abbreviations are PS, photosystems; ET, electron transport; PPP, pentose phosphate pathway; FA, fatty acid; TCA, tricarboxylic acid cycle; ETC, electron transport chain; BCAA, branched chain amino acid.

Many features characteristic of quiescence are found in *F. cylindrus* in prolonged darkness (*32*), such as the overall reduction of metabolic activity, shown by the down-regulation of RNA polymerase II subunit genes (**Figure 5)**, and by the drastic decrease in the active form of RNA polymerase II protein by immunoblot **(<u>Figure S7</u>**). On average, the total protein level per cell drops to 22% (standard deviation of 13%) after 1 month in the dark as compared to T0LIGHT (**<u>Figure S1K</u>**), consistent with the downregulation of genes involved in ribosome biogenesis in prolonged darkness (**Figure 4C**). As found in quiescent yeast, human and other animals, cell size is slightly reduced in the dark (**<u>Figure S1A</u>**) (*21*).

In prolonged darkness, the profile of expression of genes involved in glycolipid metabolism (digalactosyldiacylglycerol synthase DGDG1 and sulfoquinovosyltransferase SQD2) and in TAG synthesis (glycerol-3-phosphate acyltransferase GPAT and diacylglycerol acyltransferase DGAT) along with the decrease of thylakoid lamellae thickness suggest a potential rearrangement of cellular lipids, with thylakoid membrane lipids being reassigned to energy-rich reserve lipids (**Figure 3E, 5**). Most of the genes involved in fatty acid synthesis are downregulated in the dark while several TAG lipases and beta oxidation-related genes are induced. As suggested previously, branched chain amino acids play a key role in diatom adaptation and are potential substrates for TAG production (*33*). Here we find several genes involved in branched chain amino acid metabolism that show strong overexpression relative to T0LIGHT throughout the dark period. Although some cellular compounds can be converted to TAG in the dark, the cell eventually consumes its lipid stores in accordance with the TEM analysis (**Figure 3C, 3D**) and with the decrease in the cytometry fluorescence signal, reflecting the TAG stock in the LD (**Figure 5, <u>Table S4</u>**) (*34*, *35*).

TEM measurements suggest an accumulation of CHY after two months of darkness (**<u>Figure S3C</u>**), in contradiction to the upregulation of glucosidase genes and the downregulation of most genes involved in CHY production in darkness (**Figure 4C, 5**). Although some cells may accumulate CHY, in our opinion it is more likely that it is consumed. This is the case in limiting environments (*24*), and this carbohydrate is also known to accumulate during the day and be depleted in the dark in diatoms (*36*). While genes involved in glycolysis are globally expressed in the dark as well as when light returns, genes in the pentose phosphate pathway are upregulated primarily in the early stages of darkness (**Figure 5**), potentially required for the maintenance of carbon homeostasis.

A number of autophagy-associated genes appear to be upregulated in prolonged darkness. In addition, complexes I and II of the mitochondrial electron transport chain (alternative oxidase (AOX), NADH dehydrogenase (NDH), succinate dehydrogenase (SDHA)) are upregulated, whereas complexes III and IV (ubiquinol cytochrome c reductase (RIP1), cytochrome c oxidase (COX)) were not (**Figure 5**). This could suggest that the AOX pathway, which can play a role in cellular adaptation under different types of stress (*37*, *38*), is also involved in the dark.

The upregulation of light harvesting complex LHCx 6 and 8 and LHCz genes in prolonged darkness, in conjunction with the permanent stock of DD illustrate that as in other polar diatoms, non-photochemical quenching is a key strategy for energy dissipation upon the sudden return of light (**Figure 5**, **<u>S1G, S1H</u>**) (*21*). Therefore, photoprotection is induced rapidly after light return and then relaxed as the rETR_max_ and P^B^_max_ slowly increase to their light-acclimated values after 2-3 d. The immediate increase of rETR_max_ when light returns suggests a re-organisation of the existing photosystems rather than de novo synthesis (*18*, *21*, *39*, *40*). Interestingly, the continuous change from the transcriptomic short-term (6 h to 1 d) to longer-term response to light is consistent with the progressive increase in protein content per cell and in the observed changes in photosynthetic parameters. The return to the initial state of gene expression was observed after 2-3 d of light return, as the cell produces new photosynthetic pigments and divides (in conjunction with upregulation of cell division-related genes) (**Figure 4A, 5, <u>S1F</u>**). Of note, the nutrient deficiency on day 7 of light return had an impact on the “light-adapted” parameters that were manifested from 3d to 5d following light return, such as DES and F_V_/F_m_ values (**<u>Figure S1C</u>**).

## Discussion

Although polar diatoms are known to survive the polar night, the underlying physiological and molecular mechanisms that allow them to cope with darkness and light return are unclear. Our results are largely consistent with previous findings on common parameters measured but we also bring many novelties to the topic, expanding the number of features explored by combining photophysiological, biochemical, proteomic, microscopic and transcriptomic approaches, and this over up to 3 months of darkness. We demonstrate the resilience of the polar diatom *F. cylindrus*, with the majority of the population remaining viable in prolonged darkness (99%) as well as upon return to light (70%). We also propose explanations at the molecular level for these survival mechanisms.

In starved yeast, autophagy is intimately linked to proper entry into the quiescent state (*43*). In *Emiliana huxleyi*, starved cells thought to be quiescent are vacuolated and develop autophagosomes that engulf cytoplasmic contents and deliver them to the vacuole where hydrolytic enzymes degrade the contents, presumably providing essential building blocks for the cell (*41*). In addition to inorganic nutrients, light is an indispensable component for proliferation of photoautotrophic species. Similar vacuolar features have been observed in *P. tricornutum*, in Antarctic red algae and in chlorophytes, and may serve for material digestion in the dark (*18*, *40*, *44*). Consistent with the upregulation of genes encoding the central autophagy mechanism (autophagy-related protein ATG and the vacuolar sorting protein VPS) and with the appearance of a large vacuole (**Figure 2C, 3A, 5**), we suggest that *F. cylindrus* maintains energy homeostasis through autophagy in prolonged darkness. Autophagy can lead to programmed cell death under severe stress, but can also help diatoms to cope with nutrient shortages (*45*). In the microalgal literature, authors suggest that autophagy is involved in the degradation of LD sometimes involving their uptake into vacuoles (*42*, *46*, *47*). This is reminiscent of lipophagy in yeast where LD-containing vesicles can accumulate in the vacuole of the quiescent cell (*48*). The up-regulation of apoptosis-associated genes could entail an ongoing controlled autocatalytic death process or a quiescent stage. Based on the up-regulation of these genes from this period, we propose that this mechanism is initiated after 3d of darkness. (**<u>Figure S6</u>**). Finally, the FIB-SEM results and some TEM pictures suggest the presence of material inside the vacuoles in the dark that could be autophagic bodies (**<u>supp videos, Figure S3G</u>**).

Our data suggest an increase in ammonium concentration in the medium in the dark followed by a decrease (**Figure S1C**). This could be due to protein degradation (**<u>Figure S1K</u>**) that releases ammonium into the culture medium, which is toxic to the cell, and is then remetabolized. Interestingly, some key genes in the urea cycle are upregulated during the first hours of darkness, as are the associated proteins (*29*). Similarly, genes encoding ammonium and urea transporters appear to be expressed in the dark. This process could be involved in the conversion of N-rich degraded proteins to increase C-rich production (**<u>Table S5</u>**) (*49*). We also explored the possibility of the involvement of the dark nitrate respiration metabolic pathway, which is known to produce ammonium (*50*). Only genes encoding one of the key enzymes/functions were upregulated in the dark, but were further upregulated following light return as compared to T0LIGHT (**<u>Table S5</u>**). Furthermore, this metabolic pathway is supposed to occur in the absence of oxygen, which was not the case in our experiment.

Although counterintuitive because this metabolic pathway produces less ATP compared to the COX pathway, it appears that *F. cylindrus* upregulates the alternative AOX-dependent respiration pathway genes in prolonged darkness. Conversely, a previous proteomic study (*29*) showed undetectable ATP levels in prolonged darkness in *F. cylindrus* but also found that AOX partner enzymes, those in complexes I and II and V of the mitochondrial electron transfer chain, were upregulated in darkness. Conversely, COX-associated genes (and mitochondrial complex III partner enzyme) are induced immediately after the return to light (**Figure 5, <u>S4</u>**). Bearing in mind that our dark experiments are contrasted with T0LIGHT, it is therefore possible that respiration is mediated by COX in prolonged dark conditions, but that AOX participation is relatively more important compared with continuous light. Evocative of the case of quiescent yeast, the AOX pathway also releases fewer reactive oxygen species. Nevertheless, catalase and glutathione peroxidase genes were up-regulated in the dark and during the light return phase (**<u>Table S5</u>**). We also note that the peak in respiration is consistent with the peak in primary production after 1d of light return (**Figure 1G, 1H**).

Dark conditions of 80+ days are encountered in the Arctic between 72°N and 78°N in winter. We believe that the drastic illumination conditions at relatively high intensities applied here reflect what might occur in the field during a sudden ice breakup. We believe that similar results would have been obtained by applying a lower light level since a recent study has shown that phytoplankton growth is possible even under the ice at extremely low light intensities (*51*). Another set of samples that spent one month in the dark were also subjected to transcriptome sequencing. The positioning of the samples in the principal component analysis (**<u>Figure S5</u>)** reveals a similar delay in returning to a state of continuous light acclimation after 7d return to the light. This illustrates the resilience of polar diatoms to varying lengths of polar night that can be experienced in the natural environment.

While most of the genes involved in Chl synthesis were downregulated and the Chl a pool per cell decreased in prolonged darkness (**<u>Table S5, Figure S1F</u>**), genes encoding protochlorophyllide reductase POR and magnesium chelatase H subunit were upregulated in darkness. As previously suggested (*52*), the duplication of protochlorophyllide reductase POR genes in microalgae might allow for neo-functionalization, which here could ensure a ready supply to convert protochlorophyllide into chlorophyll when light returns (*53*). While the maintenance of magnesium chelatase H subunit gene expression in prolonged darkness may also be involved in Chl production, this particular subunit is also known to contribute to retrograde chloroplast signaling. We therefore propose that organelles send signals to the nucleus to control nuclear gene expression in the absence of light (**Figure 5**, **<u>Table S5</u>**) (*54*). Similarly, genes encoding a subunit of PSII (PsbU), enabling the essential step of water photolysis, were maintained in the dark which might promote rapid reversibility to photosynthetic metabolism when light returns (**Figure 5**).

Interestingly, DNA repair genes are maintained in tardigrades and mammals during hibernation. Here the maintenance of expression of the cryptochromes CPF2 and 3, CPD photolyase DNA repair and RAD family DNA repair genes in prolonged darkness (**Figure 5**) may contribute to the resistance of diatoms to extreme environmental conditions. Similarly, many genes encoding chitinase are upregulated in darkness, reinforcing the hypothesis of a role for this gene family in responses to environmental changes in diatoms (*55*). The persistence of the diatom-specific cyclins dsCYC2 and 3 in the dark may be involved in cell division arrest signaling driven by interaction with photoreceptors such as aureochromes, although the latter do not appear to be regulated at the transcriptomic level (*17*, *56*) (**<u>Table S5</u>**).

Finally, the observation that P^B^_max_ fell to zero after 2 months of darkness, while rETR_max_ remained above zero, suggests that some capacity for PSII activity is retained through prolonged dark but that it does not flow through to the linear chain of electrons that ultimately leads to carbon fixation mediated by RuBisCo (*21*). The residual PSII ETR upon measurement may therefore drive alternative electron transfers, perhaps involving cyclic transport around PSII (*57*).

The long-term survival strategy in the dark in *F. cylindrus* therefore appears to involve stimulation of a state of quiescence that relies on autophagy, minimal metabolic rates based on reduced RNA polymerase activity, as well as low rates of respiration that restricts production of reactive oxygen species. The overexpression in the dark of genes involved in CHY degradation, β-oxidation, and branched chain amino acid breakdown suggests that the cell feeds respiration from sugars, lipids and some amino acids, respectively. Overall, it seems that the polar diatom adopts a “better prevent than cure” strategy that is characterized by the induction of genes involved in photoprotection but also in DNA repair throughout the dark period in anticipation for the return to light. The role of the vacuole requires further study but we believe that it could be involved in the degradation of compounds by autophagy.

Although the hypothesis of an enhanced involvement of the AOX-dependent respiration in the dark compared to light conditions is likely to be relevant here, the alternative hypothesis of the maintenance of AOX gene expression in the dark in preparation for the return to light cannot be ruled out. This would require measurement of the actual activity of AOX. Furthermore, genetic/pharmacological inhibition of genes encoding pathways involved in autophagy should be explored to determine if this has a dramatic effect on entry into the quiescent state and long-term survival in the dark. The case of the maintenance of expression of the gene encoding magnesium chelatase H in the dark is also intriguing and should be explored by genetic inhibition to depict the role of retrograde signaling from chloroplasts. More detailed studies in the dark could furthermore have allowed us to determine the time of vacuole formation, its contents, and monitoring of how long cells can survive under such conditions without the consumption of constituents being detrimental to the viability and survival of the cell upon return to the light.

## Materials and Methods

### Experimental design

The cultures of *F. cylindrus* CCMP 3323 were kept in 80 Liter custom-made borosilicate glass cylinders (1.4 m tall and 30 cm in diameter) in Aquil media made from artificial sea water (*58*). Cultures were stirred with 15 cm magnetic stir bars and constantly bubbled with air (filtered through active charcoal and two 0.1μm filters Cytiva Poly-Vent™) to avoid carbon limitation and oxygen deprivation. Light was provided by a series of LED light strips (Cool white Light Reel 65 – 151 Lumens/ft) glued vertically on the two halves of a cardboard cylinder lined with tin-foil tape. Gelatin filters (LEE Filters, Hampshire UK, (#007: Pale Yellow, #138: Pale Green and #137 Special Lavender) were inserted between the LEDs and the culture to correct the light spectrum to get closer to that experienced by phytoplankton under sea ice. Light intensity was controlled by a 0-10 VDC low voltage dimmer connected to a rotary switch and adjusted daily at 30 μmol quanta m^-2^ s^-1^. The triplicate culture systems were kept inside a cold laboratory at 0 ± 1°C (humidity below dew point). Temperature of the cultures was measured daily using a thermocouple sensor.

For 6 weeks, the triplicate cultures were brought progressively up in volume following a semi-continuous protocol with 2 transfers per week. After 6 weeks, cultures were transferred in the cylinders for a final volume increase to reach 80 L. A total of 9 weeks of semi-continuous culturing at 2 transfers a week insured a total number of population generations sufficient to reach balanced growth. During the darkness period, all work in the laboratory was performed under green light (LEE filter #740) (PAR = 0.11 ± 0.03 μmol quanta m^-2^ s^-1^). Sampling was through a sampling port 10cm above the bottom of the cylinder. Growth of bacteria in the cylinders was contained by adding a combination of Streptomycin (100 mg/L), Kanamycin (100mg/L), Antimycin (100mg/L) and Gentamycin (50mg/L) to the medium before darkening (**<u>Figure S3B</u>**). Culture parameters are summarized in **<u>Table S1</u>**.

### Sample preparation and microscopy acquisition

The cultures for TEM, SEM and FIB-SEM microscopy were grown at 4°C, firstly acclimated to full light at 30 μmol quanta m^-2^ s^-1^ (light samples) and then subjected to dark for 2 months (dark samples). For sampling cells were pelleted in a cold centrifuge at 1000 rcf for 10 min. For SEM and FIB-SEM, the concentrated cells were chemically fixed in 2.5% glutaraldehyde and 4% paraformaldehyde in 0.15M marPHEM buffer (*59*) for 72h before being transferred to 0.15M PHEM containing 4% paraformaldehyde. For SEM, cells were washed 6 times with H2O, resuspended in 200 μl of H2O, and pipetted onto a 0.65 μm mixed cellulose ester mesh filter (MF-Millipore, DAWP02500) before air drying. Image acquisition was performed with a Zeiss CrossBeam 540 and imaging performed with 5 kV acceleration using a secondary electron secondary ion detector. For FIB-SEM, cells were washed 3 times with 0.15M PHEM and concentrated before being aspirated into cellulose capillaries (Wohlwend GmbH). The capillaries were cut and closed into 1-2 mm pieces and positioned in 2 mm wide and 200 μm deep aluminium carriers (Wohlwend GmbH) coated with hexadecane (Merk). Samples were cryofixed using an HPM010 high pressure freezer (AbraFluid). Freeze substitution (FS) and resin embedding were performed in an AFS2 automated machine (Leica), using the processor unit for FS. FIB-SEM acquisition was performed with a Zeiss CrossBeam 550 at the EMBL Electron Microscopy Core Facility, using the Atlas workflow (Fibics Inc.). Precise milling during the run was performed with a current of 700 pA and imaging was done with an acceleration of 1.5kV using a backscattered electron detector. An isotropic voxel size of 5 nm and a dwell time of 10 μs were used. FIB-SEM image stacks were aligned using the “Alignment to median smoothed template” workflow described by Hennies et al, 2020 (n=9 cells per condition). Segmentation was performed using MIB. Amira was used for morphometric analysis and visualization (n=3 and 5 for dark and light conditions respectively).

For TEM, cells were directly cryofixed after centrifugation, using a high-pressure freezer (HPM100, Leica) in 200 μm deep, 2 mm wide aluminum carriers (Wohlwend GmbH). Cryofixed samples were subjected to freeze substitution (EM AFS2, Leica) using the following program and buffers: 98h at −90°C in 1% tannic acid and 0.5% glutaraldehyde in acetone, 5 washes at −90°C with acetone, 32h at −90° C in 2% bone tetroxide in acetone, heating rate of 5°C/h for 14h at −20°C, 16h at −20°C, heating rate of 6°C/h for 4h at 4°C, 5 washes at 4°C with acetone. The cells were then progressively infiltrated with EPON hard resin. The resin (without accelerator)/acetone (v/v) series used were 25% for 2h starting at 4°C and reaching 20°C with a heating rate of 8°C/h, 50% for 2h at 20°C and 70% for 2h at 20°C. Infiltration with a 100% resin-containing accelerator was then performed for three times 3h and one time O/N before polymerization at 60°C for 48h. 70nm thin sections were obtained using an ultramicrotome (LeicaEM) with an ultra-diamond knife (Diatome). Two by two tiles were acquired semi automatically on the JEOL JEM 2100plus using 120Kv and 8000X magnification. Micrographs were processed using Fiji and tiles stitched using photoshop.

The cultures for confocal microscopy were grown at 4°C, firstly acclimated to full light at 30 μmol quanta m^-2^ s^-1^ and then subjected to dark for 1 month and a half (dark samples), followed by a return to 30 μmol quanta m^-2^ s^-1^ for few hours. Confocal microscopy was performed using a Leica TCS SP8 at the IBENS imaging facility (IMACHEM-IBiSA). Briefly, cells were first stained with NileRed in their culture medium (Merck - Sigma aldrich #72485-100MG) at a final concentration of 2.5 ng/mL for 20min in the dark with shaking. The cells were then centrifuged at 10,000g and the pellet resuspended in DAPI (4’,6-Diamidino-2-Phenylindole, Dihydrochloride; Thermofisher #D1306), at a final concentration of 0.1 mg/mL.

### TEM analysis and statistical analysis

After removing duplicate images, 86 Light and 99 Dark images were analyzed. All quantitative measurements were performed using Fiji 2.3.0/1.53f, with either the free-hand tool or the line tool. Heterochromatin vs euchromatin ratio was measured by binarizing the nucleus and counting the number of dark over white pixels. The nucleolus has been removed from the measurement of heterochromatin on euchromatin and has been considered independently. If the repartition of a given feature of interest followed a normal distribution (assessed with a shapiro test), means were compared with a two-sided student test considering whether the two populations’ variances were equal or not upon the result of a Fisher test. Otherwise, if the populations compared were large enough (n_i_ > 30), means were compared with a two-sided student test with populations’ variances considered unequal. Otherwise, a non-parametric two-sided Wilcoxon test was performed. All tests’ results are given with a 5% confidence level. NS stands for not significant; * stands for p-value (p) of 1% ≤ p < 5%; ** for 0.1% ≤ p < 1% and *** for p < 0.1%.

### Biochemistry and cytometry measurements

Cell number and size distribution of phytoplankton cells were determined using a Multisizer 4e (Beckman Coulter, Brea, CA, USA) coulter counter equipped with a 20μm aperture. Samples were diluted using salted Milli-Q™ water (35g/L) filtered through 3 successive 0.2 μm polycarbonate filters to avoid contamination.

Bacterial contamination, cell viability and neutral lipids content were assessed using flow cytometry (Guava easy-cyte, blue LASER at 480 nm, EMD Millipore, Darmstadt Germany). SYTOX™ Green (5nM in DMSO) dead cell stain (Invitrogen™, Thermo Fisher Scientific, MA, USA) was used to assess cell’s viability. SYBR™ Green (Invitrogen™, Thermo Fisher Scientific, MA, USA) was used in conjunction with natural chlorophyll *a* red fluorescence to assess bacterial contamination. BODIPY™ 505/515 (4,4-Difluoro-1,3,5,7-Tetramethyl-4-Bora-3a,4a-Diaza-s-Indacene; Invitrogen™, Thermo Fisher Scientific D3921, MA, USA) was used to determine the phytoplankton neutral lipid content.

Macronutrient concentrations were determined on diluted samples (salted Milli-Q™) using an autoanalyzer 3 (Seal Analytical, Wrexham, UK) segmented flow analyze. Particulate C and nitrogen content were measured by combustion using a CHN elemental analyzer (Perkin Elmer 2400 series II, Akron OH, USA). Samples were filtered onto pre-ashed (450°C, 4h) GF/F filters (Cytiva™, Marlborough MA, USA), and dried for 24h at 60°C before analysis.

Respiration rates of the cultures were assessed by measuring the change in dissolved oxygen concentration in the dark, in a culture sample using optical sensors (FireSting®-O_2_, Pyro Science GmbH, Aachen, Germany). The sample was gently stirred to avoid cell sinking during the measurement. Sampling bottles were airtight with the measuring probe applied through the stopper. Oxygen concentration was measured for 1h. Data was fitted using a simple linear model, respiration being the slope of the model (in μmol O_2_ l^-1^ h^-1^).

Concentrations of photosynthetic and non-photosynthetic pigments were determined by HPLC (*60*). Samples (10mL) were filtered onto GF/F filters (25mm), immediately flash frozen in liquid N, and stored at −80°C until analysis. After extraction in 100% methanol, samples were mixed (70/30 v/v) with a solution of tetrabutylammonium acetate (28 mM). Separation and identification of the different pigments was done on an Agilent Technologies HPLC chain equipped with a Zorbax Eclipse XDB-C8 3.5 μm column. The de-epoxidation state (DES, in %) is calculated as in (equation 1):

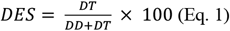

With DT is diatoxanthin and *DD* is diadinoxanthin (in mol per 100 mol chlorophyll *a*).

### Photosynthetic performances

A red light source (λ = 655 nm) pulse amplitude modulated (PAM) fluorometer (PHYTO-PAM with PHYTO-ED unit, 470nm mesuring light, 655nm actinic light, Walz, Germany) was used to measure the photosynthetic performance of *F. cylindrus* cells on dark-acclimated (30 min) samples. Rapid Light Curves (RLCs) were performed with 13 steps of 30 s and of increasing light intensity from 0 to 490 μmol quanta m^-2^ s^-1^. The dark-acclimated photochemical efficiency of photosystem II (F_V_/F_m_) was calculated as in (equation 2):

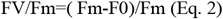

were F_0_ and F_m_ are, respectively, the minimum and maximum levels of dark acclimated chlorophyll fluorescence. The relative electron transport rate (rETR) was calculated as in (equation 3):

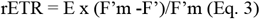

where E is the light intensity, and F’ and F_m_’ are, respectively, the minimum and maximum levels of light acclimated chlorophyll fluorescence. The determination of rETR for each of the 8 intensities of the RLCs allowed to build rETR vs. E curves (*61*) in order to extract photosynthetic parameters (*62*): rETR_max_, the maximum relative electron transport rate; *α*, the maximum light efficiency use; E_k_, the light saturation coefficient or ‘photoacclimation’ parameter; and E_opt_, the optimal light intensity for reaching rETR_max_.

The ouput of the RLCs also allowed to calculate the non-photochemical quenching (NPQ) as in (equation 4):

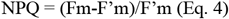

The determination of NPQ for each RLCs step allowed to build a NPQ vs E curve used to estimate the half-saturation intensity of the NPQ (E50_NPQ_) by fitting it with the model of Serôdio and Lavaud (*63*). Additionally, the maximal NPQ (NPQ_max_) was the maximal value obtained for the highest RLC step (490 μmol quanta m^-2^ s^-1^).

We repeatedly used variable fluorescence measurements to assess the culture’s photophysiological health. A triplicate of dark acclimated (30 min) sample was submitted to a blue light saturating flash (100μs, 455nm) in a Fluorescence Induction and Relaxation (FIRe) fluorometer (Satlantic, Halifax, Canada). The PSII absorption cross-section (σ_PSII_) was calculated as the initial slope of the Single Turnover fluorescence induction curve (*64*).

Photosynthetic parameters were determined from the relationship between C-fixation and irradiance by measuring the ^14^C-uptake of phytoplankton samples exposed at 24 different light levels for 20min (*28*). Raw data were fitted to the model of Platt (*65*) (equation 5):

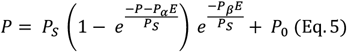

Where *E* is the irradiance of a given subsample (in μmol quanta m^-2^ s^-1^), *P_S_* is the maximum carbon fixation rate in the absence of photoinhibition (in mgC m^-3^ h^-1^), P_α_ is the light-limited slope of the carbon fixation [in mgC m^-3^ h^-1^ (μmol quanta m^-2^ s^-1^)^-1^], P_ß_ is the photoinhibition coefficient [in mgC m^-3^ h^-1^ (μmol quanta m^-2^ s^-1^)^-1^], and *P_0_* is the intercept of the curve on the y axis (in mgC m^-3^ h^-1^). The maximum rate of carbon fixation (*P_max_*, in mgC m^-3^ h^-1^), was calculated as (equation 6):

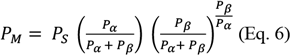

The saturation parameter, P_EK_ (in μmol quanta m^-2^ s^-1^) was calculated as *P_M_* / P_α_.

### Molecular biology procedures

Photosynthetic protein quantification was achieved by immunoquantitation (*66*). Samples (40mL) were harvested by centrifugation (4400rpm, 15 min. at 0°C) after addition of Pluronic^®^ (1μl/mL) and washed using marine PBS. Immediately after the final centrifugation, samples were flash frozen in liquid nitrogen and stored at −80°C until analysis. Proteins were extracted with 300uL of protein extraction buffer and lysed for 3 cycles of 60 seconds at 6.5 m s-1 with 60 second intervals on ice between cycles using a FastPrep-24 homogenizer. The DC™ (detergent compatible) protein assay was used for total protein determination using a calibration curve of bovine gamma globulin (BGG) standard in the 0 to 1.0 mg/mL range. Sample extracts and target standards (PsbA, Agrisera AS01 016S and RbcL, Agrisera AS01 017S) were loaded on 17 well Invitrogen Bolt 4-12% Bis-Tris gels and separated by electrophoresis using the Invitrogen Bolt Mini Gel Tank. Gels were transferred onto PVDF membranes, subsequently incubated for 1h at room temperature in their appropriate primary antibody, PsbA (Agrisera, AS05 084) or RbcL (Agrisera, AS03037). Blot membranes were imaged using a Li-Cor C DiGit Blot Scanner by chemiluminescent detection after membranes were incubated in a goat anti-rabbit IgG horseradish peroxidase secondary antibody (Agrisera AS09-602) followed by Amersham ECL Select chemiluminescent substrate. Target protein concentrations were quantitated using Image Lab 6.0.1 software (Bio-Rad).

Immunoblots were also performed from cells acclimated to full light and then subjected to one month of darkness in conditions similar to the samples for microscopy (4°C). For the RNA polymerase II, chromatin was extracted (*67*) and we use the antibody RNA pol II CTD phospho Ser2 antibody (AbFlex 91115) that binds to a post-translational modification characteristic of the active form of RNA Polymerase II. Histone H4 was used as a loading control, the gel was 6% of acrylamide and the transfert at 30V overnight. Quantification of the blot had been performed using ImageJ2 v.2.9.0. The result reflects the relative amount as a ratio of each protein band relative to the lane’s loading control using the mean grey value of the bands.

Cells for transcriptomics were pelleted in a cold centrifuge, rinsed with mPBS (15mM Na_2_HPO_4_, 4.5mM NaH_2_PO_4_, 490mM NaCl and 0.1mM EGTA) and plunged into liquid N. RNA was isolated following the protocol of the manufacturer using the RNeasy Mini Kit (Qiagen). A second DNase step was added after elution (DNase Invitrogen™ Kit TURBO DNA-free). Library preparation was made using illumina TruSeq Stranded mRNA Library Prep including poly-A selection, performed at the biotechnology company Fasteris along with sequencing that yield more than 30 million reads per sample (Illumina Sequencing HiSeq 4000 Paired-reads, 2x 150 bp).

### Reference genome and bioinformatics

The reference genome (*19*) was generated from the non-axenic monoclonal strain CCMP1102. In the case of our experiments, we used the axenic monoclonal strain CCMP3323, which is considered to be synonymous with the CCMP1102 strain. A revision of the reference genome has recently been published (*68*), suggesting a potential triploid character of this species. Despite this, we have used the genome annotation published in 2017 and believe that it is a valid source of information on the species of interest. From this reference genome, we removed genes that overlap with transposable elements (see below). These will be analyzed in a future study focusing on the expression of transposable elements during prolonged darkness. Finally, we are working with 23,902 genes that do not overlap with transposable elements.

We ran a *de novo* repeat detection on the reference genome assembly using the TEdenovo pipeline from the REPET package v2.4 (*69*) (parameters were set to consider repeats with at least 5 copies). We obtained a library of 1,115 consensus sequences that were filtered to keep only those that are found at least once as full length copy in the genome assembly and we retained 782 of them. This library of consensus sequences was then used as digital probe for whole genome annotation by the TEannot (*70*) pipeline from the REPET package v2.4.

Raw data were deposited at https://www.ncbi.nlm.nih.gov/geo/ under the GEO accession number GSE218215. The quality of raw data (150 bp paired-end reads) was verified with FastQC v0.11.9 (available at https://www.bioinformatics.babraham.ac.uk/projects/fastqc/). Adapter sequences were deleted with Trimmomatic v0.38 (*71*) with the command “trimmomatic-0.38 PE -threads 8 -phred33 -validatePairs FORWARD.fastq REVERSE.fastq ILLUMINACLIP:TruSeq3-SE.fa:2:30:10 LEADING:5 TRAILING:5 MINLEN:20”. The trimmed reads were aligned to the reference genome of *F. cylindrus* using STAR v2.7.9a (*72*) using the command “STAR -- alignIntronMax 5000 --alignIntronMin 20 --outFilterMismatchNmax 2 --outFilterMatchNminOverLread 0 -- outMultimapperOrder Random --outFilterScoreMinOverLread 0” in order to capture all the reads including the short ones. Raw read counts over exons were obtained with htseq-counts v0.13.5 (*73*) with the command “htseq-count -s reverse --nonunique all -f bam -r pos -i transcriptId -t exon -a 0 INPUT.bam Fracy1GeneModelsFilteredModels1.exons.gff” and used for differential expression analysis with DESeq2 v1.26.0 (*74*) using T0LIGHT as the reference time point. Principal component analysis was performed using DESeq2 v1.26.0. Macro-trends of gene expression were defined with the Dynamic Regulatory Events Miner, DREM 2.0 (*75*) taking in account only genes differentially expressed compared to T0LIGHT (ļlogFCļ > 2ļ in at least two consecutive time points and a penalized likelyhood node penalty value of 700. Sub-clusters of genes within each macro-trend were identified with the function kmeanspp of the package LICORS 0.2.0 (*76*).

The R package topGO was used to determine the gene ontology GO terms enriched in each sub-cluster of genes (Fisher’s exact test, p < 0.05) and used CirGO to visualize them (*77*). We further explored the gene function using the KAAS - KEGG Automatic Annotation Server (*78*) for ortholog assignment. *F. cylindrus* often has several gene accessions coding for the same enzyme/function, in which case the log2FC of genes having the same temporal trend were averaged in **Figure 5**. One example is the NADH-dehydrogenase (NDH) function, for which some variants are expressed in the dark but others seem to take over when the light returns (**Figure 5**). For readability purposes, we did not include all variants nor all functions, the ones included in **Figure 5** are in **<u>Table S4</u>** and the full list of genes manually annotated is available in **<u>Table S5</u>**.

In silico targeting predictions were performed for all protein sequences using HECTAR (*79*) and ASAFind v2.0 (*80*). In fine, we illustrated photosynthesis and fatty acid synthesis in the chloroplast, beta oxidation, TCA cycle, mitochondrial ETC and branched chain amino acid metabolism in mitochondria, glycerolipid metabolism between chloroplast and endoplasmic reticulum, while glycolysis is illustrated in the cytosol (**Figure 5**).

## Acknowledgments

We acknowledge the contributions of the non-authors of this paper, including all staff who participated in the numerous sampling events. We thank Leila Tirichine for her contribution in the early stages of this project. We thank Natalie Donaher and Mireille Savoie for analyzing the protein data and Gabrièle Deslongchamps for the nutrient data. We thank Astou Tangara, Fatima Melouki and Benjamin Mathieu from the IBENS imaging facility (IMACHEM-IBiSA), a member of the France-BioImaging National Research Infrastructure (ANR-10-INBS-04), which received support from the “Federation for Brain Research - Rotary International France” (2011) and from the “Investissements d’Avenir” ANR-10-LABX-54 MEMOLIFE program, for their help in acquiring the confocal microscopy images. We also thank Florian Maumus for his kind detection and annotation of the transposable elements. Finally, we thank the prestigious EMBL Electron Microscopy Core Facility where the electron microscopy was performed.

## Author contributions

Conceptualization: MB, ChB, NJ
Methodology: NJ, LC, KM, JG, JL, SG, TS, FB, MBé, MHF, TL, JET, DC, JoL, YS
Investigation: NJ, LC, JG, JL, SG, FB, OAM, ClB, DG
Visualization: NJ, LC, KM
Supervision: MB, ChB
Writing—original draft: NJ
Writing—review & editing: all co-authors

## Data and materials availability

All data are available in the main text or the supplementary materials.

