## Supplementary figures for "Hypometabolism to survive the long polar night in the diatom *Fragilariopsis cylindrus*"

Science AAAS

**This PDF file includes:**

Figs. S1 to S7
Captions for Movies S1 to S6

**Other Supplementary Materials for this manuscript include the following:**

Table S1 to S5 [Joli-et-al-Supplementary-Tables.xlsx]
Annotation file for gene [Joli-et-al-GENES.gff]
Annotation file for transposable elements [Joli-et-al-TE.gff]

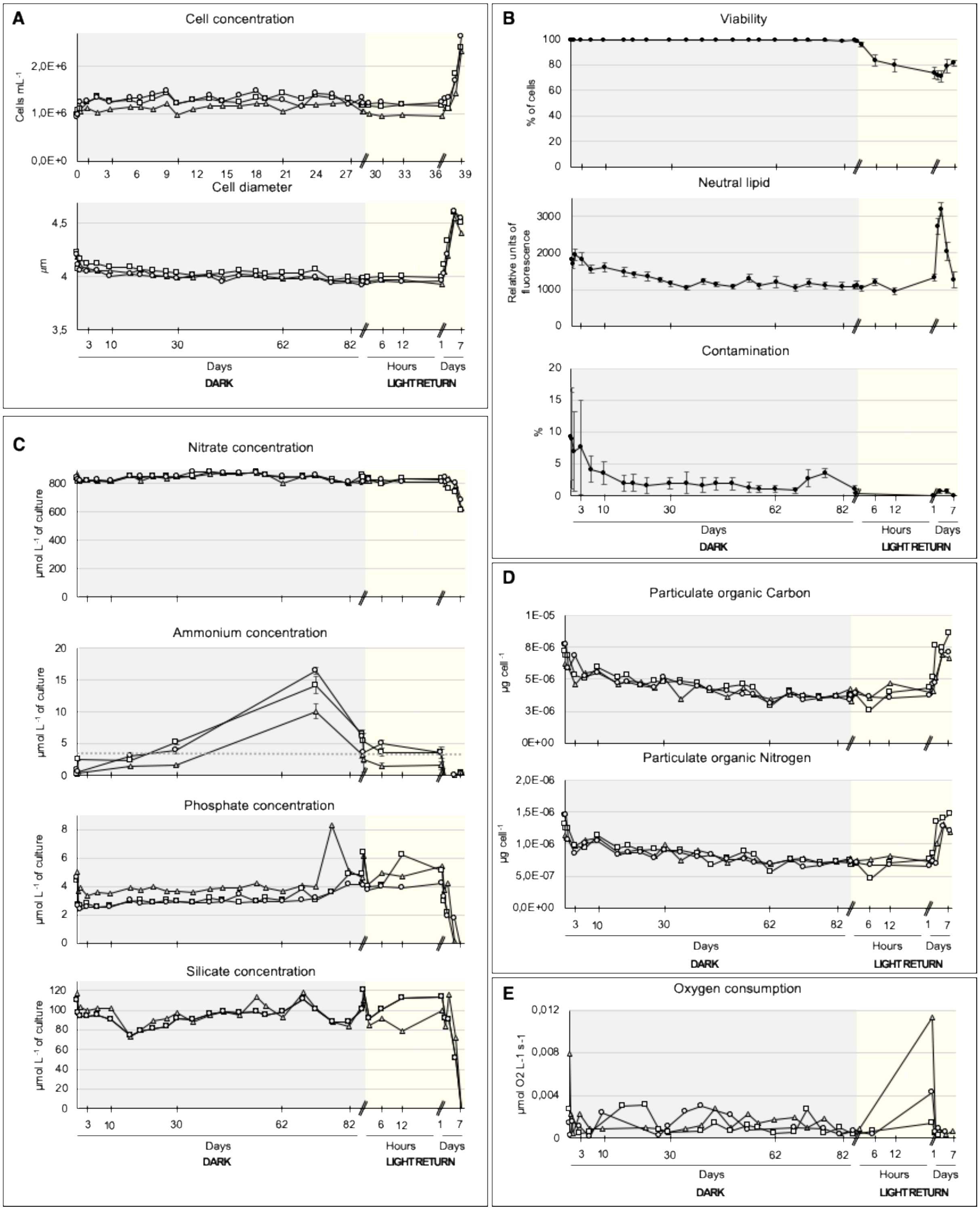
A
Cell concentration
Cells mL-1
2,0E+6
1,0E+6
0,0E+0
0 3 6 9 12 15 18 21 24 27 30 33 36 39
Cell diameter
µm
4,5
4
3,5
3 10 30 62 82 6 12 1 7
Days
Hours
Days
DARK
LIGHT RETURN
B
Viability
% of cells
100 80 60 40 20 0
Neutral lipid
Relative units of fluorescence
3000 2000 1000 0
Contamination
%
20 15 10 5 0
3 10 30 62 82 6 12 1 7
Days
Hours
Days
DARK
LIGHT RETURN
C
Nitrate concentration
µmol L-1 of culture
800 600 400 200 0
Ammonium concentration
µmol L-1 of culture
20 15 10 5 0
Phosphate concentration
µmol L-1 of culture
8 6 4 2 0
Silicate concentration
µmol L-1 of culture
120 100 80 60 40 20 0
3 10 30 62 82 6 12 1 7
Days
Hours
Days
DARK
LIGHT RETURN
D
Particulate organic Carbon
µg cell-1
1E-05 8E-06 5E-06 3E-06 0E+00
Particulate organic Nitrogen
µg cell-1
2,0E-06 1,5E-06 1,0E-06 5,0E-07 0,0E+00
3 10 30 62 82 6 12 1 7
Days
Hours
Days
DARK
LIGHT RETURN
E
Oxygen consumption
µmol O2 L-1 s-1
0,012 0,008 0,004 0
3 10 30 62 82 6 12 1 7
Days
Hours
Days
DARK
LIGHT RETURN

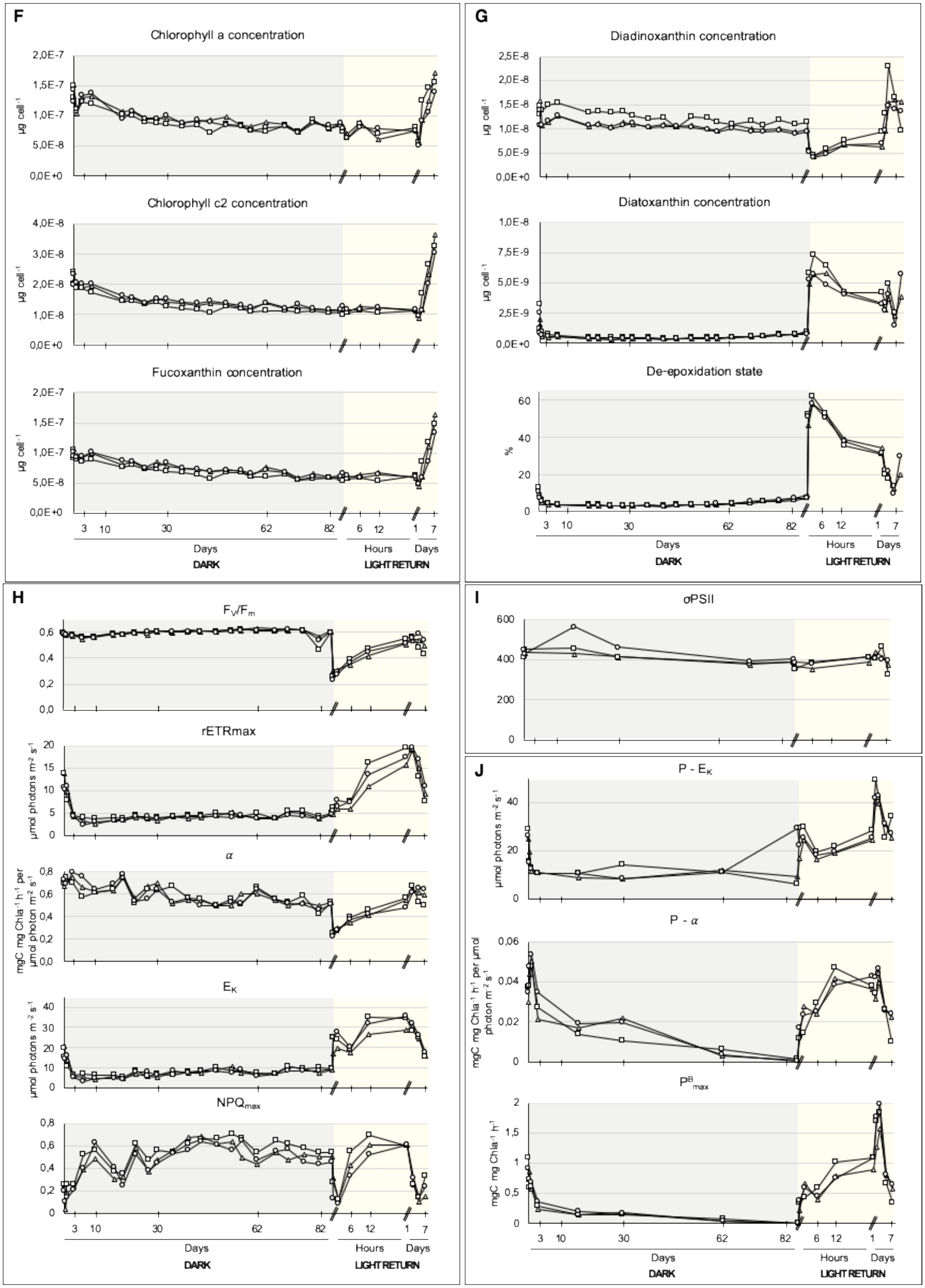
F
Chlorophyll a concentration
µg cell-1
2,0E-7
1,5E-7
1,0E-7
5,0E-8
0,0E+0
Chlorophyll c2 concentration
4,0E-8
3,0E-8
2,0E-8
1,0E-8
0,0E+0
Fucoxanthin concentration
3 10 30 62 82 6 12 1 7
Days Hours Days
DARK LIGHT RETURN
G
Diadinoxanthin concentration
2,5E-8
2,0E-8
1,5E-8
1,0E-8
5,0E-9
0,0E+0
Diatoxanthin concentration
1,0E-8
7,5E-9
5,0E-9
2,5E-9
De-epoxidation state
%
60
40
20
0
H
FV/Fm
0,6
0,4
0,2
0,0
rETRmax
µmol photons m-2 s-1
20
15
10
5
α
mgC mg Chla-1 h-1 per µmol photon m-2 s-1
0,8
EK
40
30
NPQmax
I
σPSII
600
400
200
J
P - EK
P - α
mgC mg Chla-1 h-1 per µmol photon m-2 s-1
0,06
0,04
0,02
PBmax
mgC mg Chla-1 h-1
2
1,5
1
0,5

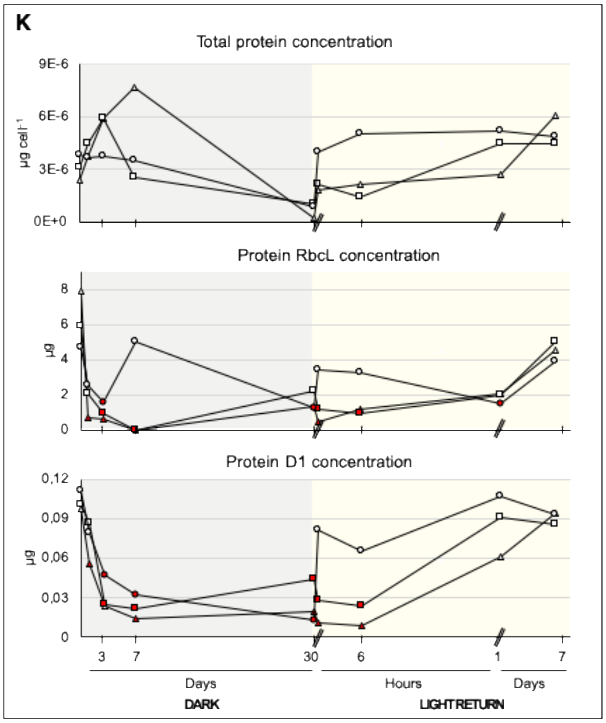


**<u>Figure S1</u>**: Results from the physiological, biochemical, cytometry and protein analysis. The triangles represent cylinder 1, squares for Cylinder 2 and circles for Cylinder 3.

**(A)** Cell abundance and cell diameter.

**(B)** Average and standard deviation of parameters measured by cytometry: percentage (%) of viability of cell (SYTOX), neutral lipid content expressed as relative units (BODIPY) and % of bacterial contamination (SYBR GREEN).

**(C)** Nitrate, Phosphate and Silicate concentrations in µmol $L^{-1}$ of culture measured with autoanalyzer. Ammonium concentration in µmol $L^{-1}$ of culture (and standard deviation) measured by incubation and fluorescence.

**(D)** Carbon and Nitrogen concentrations in µg $cell^{-1}$.

**(E)** Oxygen consumption in the culture in µmol $O_2$ $L^{-1}$ $s^{-1}$.

**(F)** Photosynthetic pigments (Chl a, Chl c2 and Fucoxanthin) concentrations in µg $cell^{-1}$.

**(G)** Photoprotective pigments (DD and DT) concentrations in µg $cell^{-1}$. The de-epoxidation state of xanthophyll cycle pigments in %.

**(H)** Parameters from Phyto-PAM Rapid Light Curves protocol; maximum quantum yield efficiency of PSII ($F_V/F_m$), maximum relative photosynthetic electron transport rate in µmol photons $m^{-2}$ $s^{-1}$ ($rETR_{max}$), chlorophyll specific photosynthetic efficiency coefficient in mgC mg $Chla^{-1}$ $h^{-1}$ per µmol photon $m^{-2}$ $s^{-1}$ ($\alpha$), light saturation coefficient in µmol photons $m^{-2}$ $s^{-1}$ ($E_K$), and non-photochemical quenching ($NPQ_{max}$).

**(I)** Average absorption cross-section σPSII measured by a mini fluorescence induction and relaxation (FIRe) in $angstrom^2$.

**(J)** Parameters from PvsE curves ($^{14}C$ incubation);  Light saturation coefficient in µmol photons $m^{-2}$ $s^{-1}$ (P - $E_K$), chlorophyll specific photosynthetic efficiency coefficient in mgC mg $Chla^{-1}$ $h^{-1}$ per µmol photon $m^{-2}$ $s^{-1}$ (P - $\alpha$) and maximum specific carbon fixation rate in mgC mg $Chla^{-1}$ $h^{-1}$ ($P^B_{max}$).

**(K)** Total protein measured in µg $cell^{-1}$. The photosynthetic proteins RbcL and PsbA concentrations are given in µg. Samples in red fall below the quantification limit.

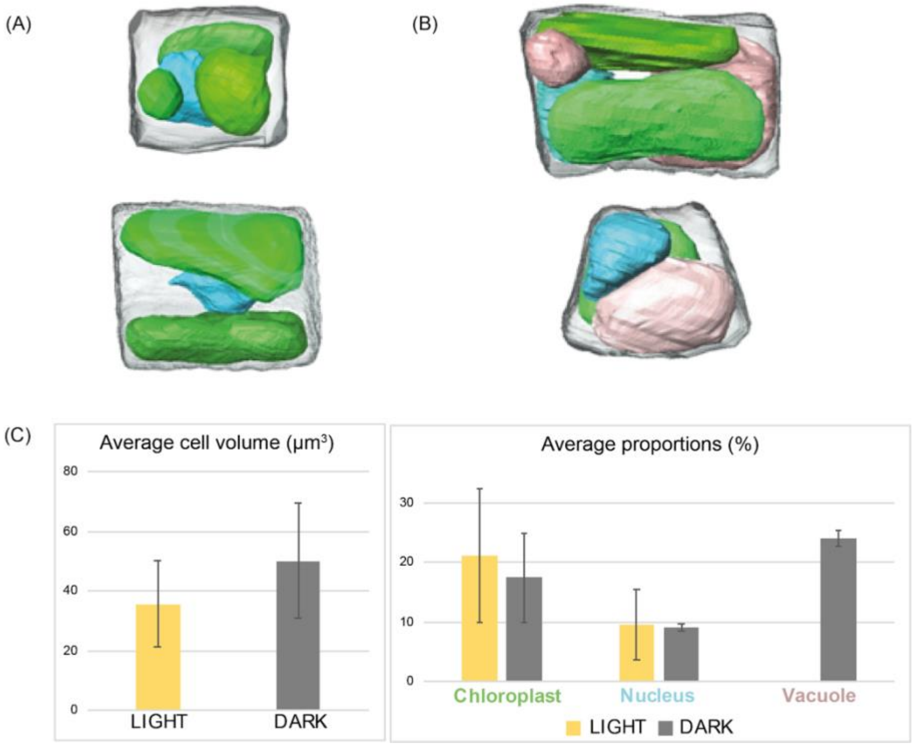


**Figure S2**: 3D Reconstitution of the cells using Focused Ion Beam-Scanning Electron (FIB-SEM) microscopy.
**(A)** Example of two light-grown cells.
**(B)** Example of two dark-adapted cells.
**(C)** Average cell volume for light (n=5) and dark (n=3) cells and average proportions of the chloroplasts (in green), nucleus (in blue) and vacuole (in pink) expressed in percentage (%) of the cell volume.

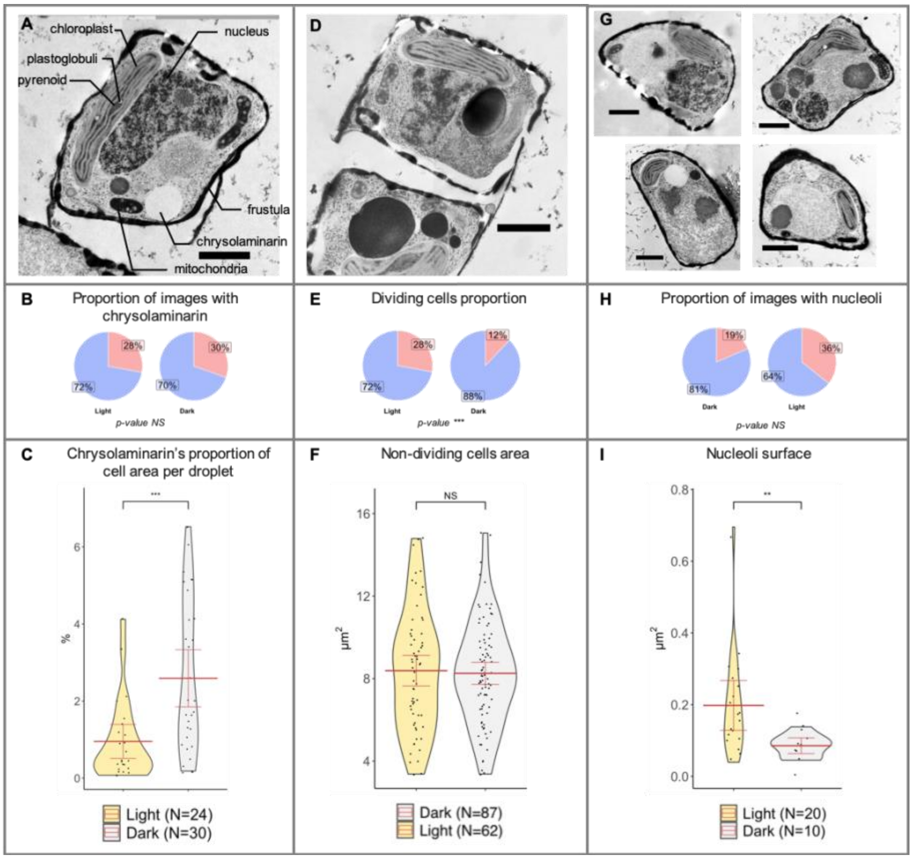


**<u>Figure S3</u>**: TEM image analysis of *F. cylindrus* in light/dark conditions show distinctive cell structural features

**(A)** Image of a dark cell illustrating the identification of the major organelle. The droplets filled with a whitish material were different from the content of vacuoles and from artifacts created during cell cutting and were considered to be CHY

**(B)** Diagram comparing the proportion of light/dark cells that do not contain chrysolaminarin (blue) to those that do (red)

**(C)** Violin plot of the proportion of chrysolaminarin per light/dark cell area per droplet

**(D)** Image of a light cell that have recently divided

**(E)** Diagram comparing the proportion of non-dividing light/dark cells (blue) to the dividing ones (red). Of note few cases of cell division were observed in the dark.

**(F)** Violin plot comparing the cells area in light/dark conditions

**(G)** Images of dark cells illustrating the presence of bodies inside the vacuoles.

**(H)** Diagram comparing the proportion of light/dark cells that do not contain a nucleoli (blue) to one that do (red)

**(I)** Violin plot comparing the surface of the nucleoli in light/dark conditions

For violins plots, mean is shown by the bold horizontal bar, its 95% confidence interval is between the two error bars. Result of mean comparison is represented by its p-value above the plots. N is the number of images used for each plot. For each image, scale bar represent 1µm

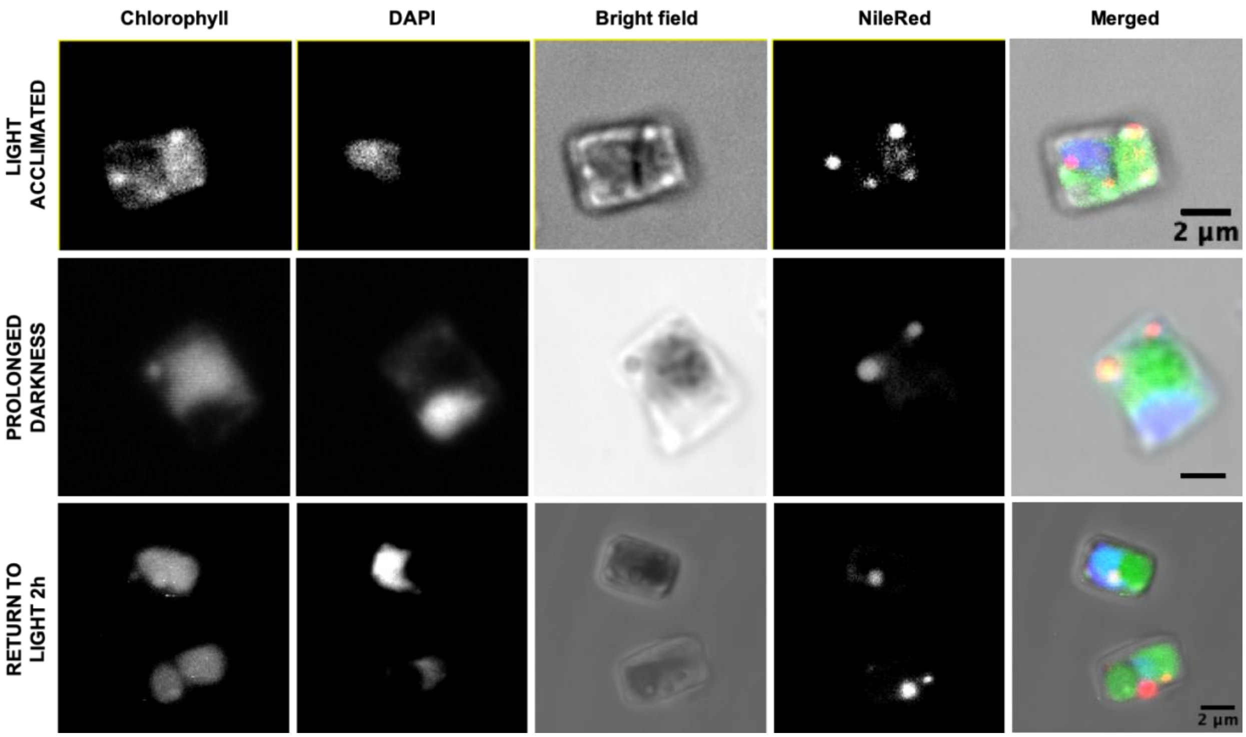


**<u>Figure S4</u>**: Fluorescence microscope images of *F. cylindrus* stained with DAPI and NileRed. Scale bar = 2 µm. Chlorophyll autofluorescence, fluorescence of DAPI, and fluorescence of NileRed are visualized in green, blue, and red colors, respectively.

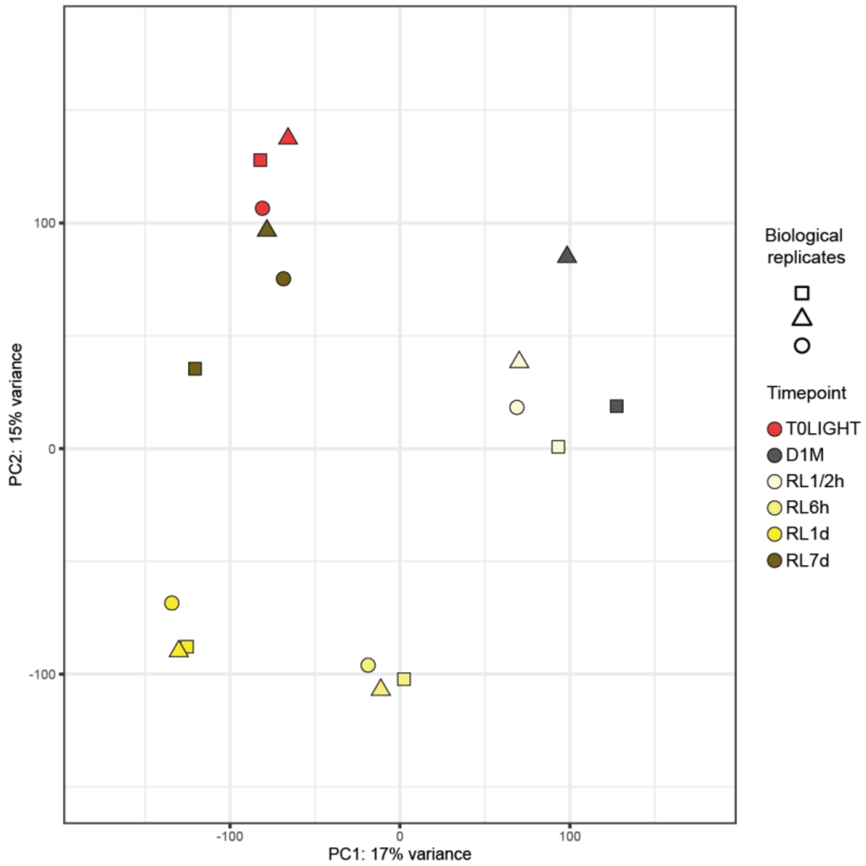


**Figure S5**: Principal component analysis of a total 17 light/dark transcriptomes from cultures in the acclimation phase in full light (T0LIGHT), in prolonged darkness (1 month), and upon return to light (up to 7 days). Each dot represents one transcriptome and the shapes refer to one of the three biological replicates.

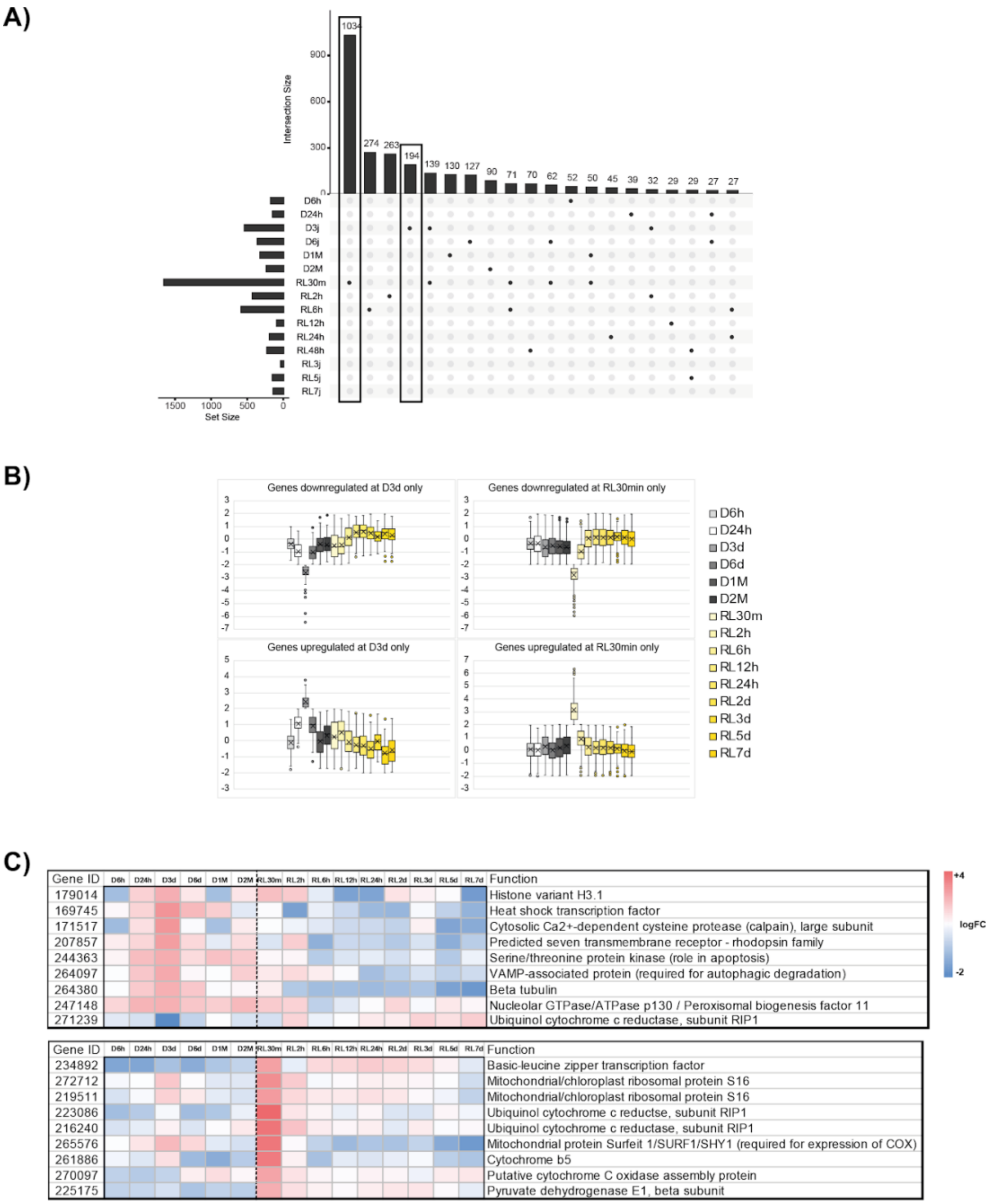


**<u>Figure S6</u>**: Tracking genes significantly differentially expressed at single time points.
**(A)** Upset plot of the 3619 genes not considered in the DREM analysis, that are significantly differentially regulated (|log2FC| > 2) in non-adjacent time points.
**(B)** Boxplot of the genes differentially expressed (|log2FC| > 2) only after 3 days of darkness (left) or after 30 minutes of light return (right). The genes had been divided in two groups based on the positive (upper panels) or negative (lower panels) log2FC compared to T0LIGHT.
**(C)** Heatmaps of selected examples of genes differentially expressed only after 3 days of darkness (upper panel) or after 30 minutes of light return (lower panel) and their biological function.

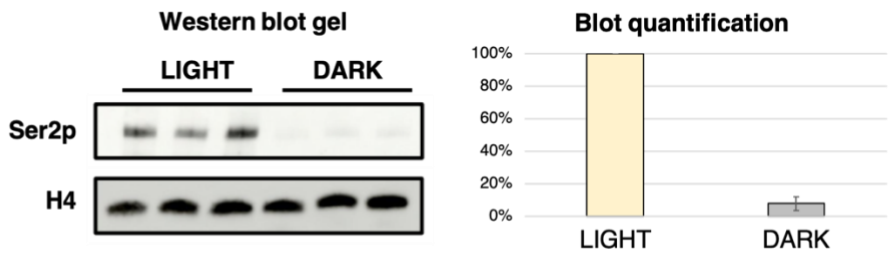


**<u>Figure S7</u>**: Western blot targeting the active form of RNA Polymerase II (RPB1 CTD Ser2p). Histone H4 was used as a loading control. On the left the gel in which the first three columns represent light samples and the last three represent dark samples. On the right the quantification of the blot, taking into account the normalization using the H4 protein.

**Movie S1.**

FIB-SEM-Light-1: 3D reconstruction of a full light-acclimated cell allowed with focused ion beam/scanning electron microscopy.
FIB-SEM-Light-2: 3D reconstruction of a full light-acclimated cell allowed with focused ion beam/scanning electron microscopy.
FIB-SEM-Light-3: 3D reconstruction of a full light-acclimated cell allowed with focused ion beam/scanning electron microscopy.
FIB-SEM-Dark-1: 3D reconstruction of a dark-acclimated cell allowed with focused ion beam/scanning electron microscopy.
FIB-SEM-Dark-2: 3D reconstruction of a dark-acclimated cell allowed with focused ion beam/scanning electron microscopy.
FIB-SEM-Dark-3: 3D reconstruction of a dark-acclimated cell allowed with focused ion beam/scanning electron microscopy.

**Data S1. (separate file)**

Data S1 to S5 [Supplementary-Tables.xlsx]

**Table S1**: Summary and timing of the parameters measured from the cultures of Fragilariopis cylindrus. All the parameters had been sampled from the same 3 biological replicates (cylinders of 80L) that had spent 3 months in the dark. * Of note, RNA polymerase II and photosynthetic protein western blots were done on samples that have spent a month in the dark and the microscopy on samples that have spent 2 months in the dark.

**Table S2**: ID of genes included within each DREM subclusters and associated enriched GO categories.

**Table S3**: Complete list of genes that are significantly differentially expressed at single time points as compared to T0LIGHT.

**Table S4**: Correspondence of genes names with acronyms, the number of genes considered and the subcellular prediction from Figure 5.

**Table S5**: Complete list of genes explored

**Annotation file** for gene [Joli-et-al-GENES.gff]

**Annotation file** for transposable elements [Joli-et-al-TE.gff]
